# Can DNA help trace the local trade of pangolins? A genetic assessment of white-bellied pangolins from the Dahomey Gap (West Africa)

**DOI:** 10.1101/2021.10.07.463484

**Authors:** Stanislas Zanvo, Sylvestre C.A.M. Djagoun, Akomian F. Azihou, Bruno Djossa, Komlan Afiademanyo, Ayodedji Olayemi, Clément Agbangla, Brice Sinsin, Philippe Gaubert

## Abstract

We conducted in the Dahomey Gap (DG) a pioneer study on the genetic tracing of the African pangolin trade. We sequenced and genotyped 189 white-bellied pangolins from 18 forests and 12 wildlife markets using one mitochondrial fragment and 20 microsatellites loci. Tree-based assignment procedure showed the ‘endemicity’ of the pangolin trade, as strictly fed by the lineage endemic to the DG (DGL). DGL populations were characterized by low levels of genetic diversity, an overall absence of equilibrium, inbreeding depression and lack of geographic structure. We identified a 92-98% decline in DGL effective population size 200-500 ya –concomitant with major political transformations along the ‘Slave Coast’– leading to contemporaneous estimates inferior to minimum viable population size. Genetic tracing suggested that wildlife markets from the DG sourced through the entire DGL range. Our loci provided the necessary power to distinguish among all the genotyped pangolins, tracing the dispatch of same individuals on the markets and within local communities. We developed an approach combining rarefaction analysis of private allele frequencies and cross-validation with observed data that could trace five traded pangolins to their forest origin, c. 200-300 km away from the markets. Although the genetic toolkit that we designed from traditional markers can prove helpful to trace the pangolin trade, our tracing ability was limited by the lack of population structure within DGL. Given the deleterious combination of genetic, demographic and trade-related factors affecting DGL populations, the conservation status of white-bellied pangolins in the DG should be urgently re-evaluated.

## Introduction

Pangolins (Order Pholidota) are considered the most trafficked mammals in the world, with c. 900,000 individuals seized over the last 20 years [1]. Although pangolins have been – mistakingly– highlighted as potential intermediary hosts of the COVID-19 pandemia [2], the volumes traded have remained unsustainably high [3]. As the exponential demand from the Traditional Chinese Medecine (TCM) market has decimated the populations of Asian pangolins, new routes of trafficking have emerged from Africa [4]. Between 2015 and 2019, an estimated number of > 400,000 African pangolins were seized en route to Asian markets [1, 5]. Consequently, African pangolins are currently experiencing unprecedented levels of harvesting, for both local demands and the illegal international trade, with a possible influence of the Chinese diaspora on the African trade networks and dynamics [4, 6, 7].

The white-bellied pangolin (WBP; *Phataginus tricuspis*) is the species with the largest range in tropical Africa and is the pangolin most frequently sold on the African bushmeat market stalls [5]. It is also the most represented African species in international seizures of pangolin scales [4, 8]. Recent genetic investigations have shown that WBP consisted of six cryptic, geographically traceable lineages [9], one of which occurs ‘outside’ the rainforest blocks, in a West African savannah corridor interspersed with very fragmented forest cover, the Dahomey Gap[10]. The Dahomey Gap lineage (DGL) is of peculiar patrimonial importance as it is endemic to a unique biogeographical zone in western Africa (from Togo to Benin and southwestern Nigeria; [9] and likely is the only pangolin species surviving in the Dahomey Gap [11, 12].

DGL populations currently suffer from intense levels of deforestation and hunting. Populations are fragmented into –generally small– patches of forest islands, and have drastically decreased in abundance through the last decades [11, 12]. In Togo and Benin, WBG is hunted for its meat and use in traditional medicine, both contributing to its overexploitation [11, 12]. Because of the recent seizure of scales intended to the international trade (Cotonou airport; [4]) the prominent proportion of DGL in Asian seizures [8] and the established traffick hub in the neighbouring Nigeria [13], there is a serious risk that DGL populations are exploited in unsustainable volumes.

The genetic toolkit has been successfully applied to trace the illegal wildlife trade, by providing accurate information on the species traded and their geographic origins [14, 15]. Recently, the genetic distinctiveness among the different WBP lineages has been used to assign the regional origins of international seizures of pangolin scales [8]. However, the current lack of knowledge on the fine-scale population structure of WBP hampers any attempts at tracing their local –e.g., country-scale– trade (see [16]). The stakes behind tracing the local WBP trade are high, as they reside in, and, therefore, better Efficient conservation actions to mitigate the local WBP trade will rely on identifying the source populations feeding the market (and thus the market network) and accurate estimates of the number of individuals traded (e.g., from scale seizures). We propose a pioneer investigation on the utility of the genetic toolkit in tracing the local pangolin trade, based on microsatellite markers recently developed for WBP [17]. Our general objective is to provide a detailed overview of the genetic status of DGL populations and their traceability on the local pangolin trade. Our specific objectives encompass the assessment of (i) population structure and diversity within DGL, (ii) the demographic history of this endemic lineage, and (iii) the resolutive power of our genetic markers to trace the pangolin trade in the Dahomey Gap.

## Material and Methods

### Genetic sampling and wet laboratory procedures

We collected a total of 189 WBP samples across Benin (6°10’ - 11°00’ N), Togo (8°10 - 9°0’) and southwestern Nigeria (6°10’ - 11°00’ N). Our sampling effort included all the occurrence zones of the species in Benin [11] and one central forest in Togo (111 samples from 18 forests), nine major traditional medicine markets (TMMs; 71) from Benin and Togo, and three bushmeat markets in Benin and Nigeria (7). Sample types varied from fresh tissue, skin and tongue (59), to dried skin and tissue (60), and scale connective tissue (70) (see Table S1, Supplementary information). Thirty-six samples were taken from carcasses having received preservartive chemical treatments [7]. Free consent from hunters and market sellers was obtained before collecting samples. We relied on an opportunistic sampling strategy [18], without financial incentives. The samples collected from the forest were traced to their original location after the information provided by hunters [11].

DNA extraction from fresh and scale connective tissues was performed using the NucleoSpin^®^ Tissue Kit (Macherey-Nagel, Hoerdt, France), following manufacturer’s recommendations. Final elution step was repeated twice in 50 µl Elution buffer to increase DNA yield. The samples treated with chemicals were extracted following a modified CTAB protocole including upstream TE washing baths and Dithiothreitol (DTT; [19]. Elution volumes varied from 30 to 100 µl nuclease free water, depending on the size of the DNA pellet. DNA concentrations were estimated on the NanoDrop 1000 Spectrophotometer (ThermoFisher Scientific, Illkirch-Graffenstaden, France).

We amplified a mitochondrial fragment of 432 bp from Control region 1 (CR1) following Gaubert et al. [9]. PCR products were sequenced at Genoscreen (https://www.genoscreen.fr/en; Lille, France) and Macrogen Europe (https://dna.macrogen-europe.com/en; Amsterdam, the Netherlands). Sequences were aligned by-eye with BioEdit v7.0.5 [20] and the unique haplotypes were submitted to Genbank under accession numbers OK275650-OK275662.

We amplified 20 microsatellite markers developed from the genome of WBP in four multiplexes following Aguillon et al. [17]. PCR triplicates were conducted for dried and chemically treated samples to mitigate the potential issue of allelic dropout and false alleles [21]. Consensus was considered met when at least two out of the three replicates indicated the presence of an allele. PCR products were separated on automated sequencer at Genoscreen and GeT-PlaGe (https://get.genotoul.fr/; INRAE, Toulouse, France).

## Data analysis – Control region 1

### Clustering

We assessed the ‘endemicity’ of the pangolins from Togo, Benin and Nigeria (N = 168) through a distance-tree analysis including all the CR1 sequences of WBP available in Genbank (N = 100). All the sequences were aligned by-eye with BioEdit v7.0.5 [20]. Phylogenetic tree reconstruction was performed in MEGA-X v10.2.2 [22] using Neightbor Joining, 1,000 bootstrap replicates, Kimura 2-parameter model [23] and Gamma distribution (G). A pangolin was considered endemic to the DGL if its sequence clustered within the Dahomey Gap lineage as defined by [9].

### Genetic diversity and structure

Genetic diversity and structure in DGL were estimated from sequences without missing data (N = 126). We used DnaSP v6.12 [24] to compute haplotype number (*h*), haplotype diversity (*Hd*) and nucleotide diversity (*π*) (Table S2, Supplementary information) for the six WBP lineages. We mapped the distribution of haplotypes in DGL using ArcGIS 10.1 (Esri France). We used Network v10.2.0.0 to build a median-joining haplotype network with ε = 0 to minimize alternative median networks.

### Demographic history

Mismatch analysis was performed in Arlequin 3.5.2 [25] to test for signatures of demographic and spatial expansion in DGL, by calculating the sum of squared deviations (SSD) between observed and expected distributions using 1,000 boostrap replicates [26].

We also tested for deviation from neutrality by computing a series of statistics in DnaSP, including Tajima’s D [27], Fu’s Fs [28], Harpending raggedness index *r* [29] and Ramos-Onsins and Rozas’ R2 [30]. We ran 1,000 replicates assuming a coalescent process with a neutral, infinite-sites model and large constant population sizes [31], to calculate the P-value of each observed statistics.

## Data analysis – Microsatellites

### Genetic diversity

Genious 9.0.5 [32] was used for allele scoring and genotype extraction through the Microsatellites plugin (https://www.geneious.com/features/microsatellite-genotyping/). Only the DGL individuals with at least 75% of genotyping success (≥15 microsatellites markers) were considered for the analyses (N = 169). Genetic diversity at each locus was characterized through (i) number of alleles (Na), observed (*Ho*) and expected (*He*) heterozysity as computed from GenAlEx 6.5 [33], (ii) allelic richness (A_R_) and *F*_*IS*_ as estimated from FSTAT 2.9.4 [34]. Deviation from Hardy-Weinberg Equilibrum (HWE) was calculated for each locus in GenAlEx. Linkage Deseliquilibrum (LD) for all the pairs of loci was tested in FSTAT using 1000 randomisations with Bonferroni correction. Null allele detection, assuming population at equilibrium, was done with Microcheker 2.2.3 [35] using Bonferroni correction.

### Genetic structure

Global genetic variance within DGL was visualized through Principal Coordinates Analysis (PCoA), using pairwise population matrix unbiased genetic distances in GenAlEx.

Pairwise differentiation (*F*_*ST*_) among forest populations (N = 104) was computed in Arlequin 3.5 [36] using three partition schemes (Fig. S3, Supplementary information): (i) a 6-partition scheme considering populations (i.e., from a same habitat patch) with ≥7 sampled individuals; (ii) a 3-partition scheme including a southern forest block radiating from the Lama forest (c. 50 km maximal radius), a central forest block radiating from the Mont Kouffé forest (c. 45 km maximal radius), and a central forest block in Togo radiating from the Assoukoko forest (c. 25 km maximal radius); and (iii) a 3-partition scheme using a latidunal-based grouping in Benin (South, lower Centre, and upper Centre). Because the first partition scheme yielded the highest levels of differentiation among populations (see Results), we calculated their inbreeding coefficients (*F*_*IS*_) in FSTAT.

We used STRUCTURE 2.3.4 [37] to conduct a clustering analysis on all the DGL individuals. We performed 20 independent runs for K = 1–10 using 10^5^ Markov chain Monte Carlo (MCMC) iterations and burnin = 10^4^, assuming admixture model and correlated allele frequencies. STRUCTURE HARVESTER 0.6.94 [38] was used to detect the most likely number of populations (K) using the □K method [39]. We also ran STRUCTURE using the LOCPRIOR model (and same other parameters) to assess the amount of information carried by the geographic distribution of populations (r). For r values > 1, the geographic information is considered uninformative [37].

We also inferred the number of populations, spatial locations of genetic discontinuities and population membership among georeferenced individuals using the *Geneland* package [40] in R 4.0.5 (*R Team Development Core 2021*). Following Coulon et al. (2006), we first allowed K to vary from 1 to 10 and launched five runs of 5.10^5^ MCMC iterations (500 thinning and 500 burn-in) under a frequence-correlated model and 1 km of uncertainty for spatial coordinates. Second, we fixed the number of estimated populations on the basis of the first analysis (K = 6-7), to perform 20 independant runs using the same parameters. We also performed 20 independent runs fixing K = 3 as obtained with STRUCTURE (see Results). For both analyses, we assessed how stable was population assignment of individuals among the best (i.e. with highest posterior probabilities) five runs. We used 500 × 500 pixels to map the posterior probabilities of population membership.

Given the absence of any clear genetic structuring from the above analyses (see Results), we performed spatial Principal Component Analysis (sPCA) using the Delaunay triangulation connection network, which defines neighbouring entities based on pairwise geographic distances. sPCA is a spatially explicit multivariate approach capable of investigating complex and cryptic spatial patterns of genetic variability from allelic frequencies[41]. Such an approach does not require data to meet Hardy–Weinberg or linkage equilibrium among loci. sPCA was run with the DGL georeferenced individuals using the *adegenet* package [42] in R, with 9,999 MCMC resampling to infer global *vs*. local structuration levels. Threshold distance between any two neighbors was set to zero.

We tested isolation-by-distance (IBD) among (i) forest individuals and (ii) populations (6-partition scheme; see above) by running a Mantel test in the R package *pegas* [43], where we quantified the correlation (*r*) between genetic (Edward’s) and geographic (Euclidean) distances through 10,000 permutations. The geographic distances among populations were calculated from the center of each forest.

### Ability of the microtellites data to trace the pangolin trade

The discriminating power of our microsatellite markers among market and non-market individuals was evaluated by (i) couting the number of identical genotypes among samples with the Multilocus tagging option in GenAlEx (suboption Matches), (ii) computing the probability of encoutering the same genotype more than once by chance using the R package *poppr* (method=single; [44], and (iii) calculating values of unbiased probability of identity and probability of identity among siblings (uPI and PIsibs) in Gimlet 1.3.3 [45].

We used the generalized rarefaction approach implemented in ADZE [46] to compute private allele frequencies among various combinations of populations. We delineated six original forest populations after partition scheme (i), and let population number vary from 2 to 6 (sample size rarefied from 2 to 7). Because the original scheme of six populations yielded the greatest frequencies of inferred private alleles (see Results), we graphed the rarefaction curves for each locus in these six populations to assess their respective contributions in the identification of private alleles per population. Loci that reached a plateau or showed an exponential trend of their estimated private allele frequencies were considered as potentially useful for tracing the origin of pangolins found in the markets, whereas loci showing a decreasing trend were discarded. We then crossed these results with the private alleles actually observed for the six populations (GenAlEx), and only considered the loci that showed both observed private alleles (GenAlEx output) and high potential for tracing (ADZE output). Finally, we manually screened the genotypes of the market individuals to retrieve such private alleles and attribute source populations.

### Demographic history

We tested for the signature of bottleneck events in DGL using the Single Mutation Model (SMM) and the Two Phase Model (TPM) in BOTTLENECK 1.2.02 [47]. We applied the Wilcoxon sign-rank test to analyze the presence of heterozygote excess/deficit using 10,000 replications.

Demographic history was also assessed through the R package *varEff* [48], an approximate-likelihood method that infers temporal changes in effective population size. Given the lack of data on sexual maturity in WBP, we fixed a conservative generation time of 2 yrs based on estimates from Asian species [49, 50]. Mutation rate was fixed to 5.10^−4^ based on published average mutation rate [51]. The analysis was run with the single mutation, geometric mutation and two phases mutation models, using 10,000 MCMC batches with a length of 1 thinned every 100 batches and JMAX = 3. The first 10,000 batches were discarded as part of the burn-in period. Confidence intervals for ancestral and current effective population size estimates were calculated from the harmonic means for each mutation model.

## Results

### Mitochondrial DNA

Our ML tree based on 268 mitochondrial DNA (mtDNA) sequences recovered the six WBP geographic lineages with robust node supports, including Western Africa, Ghana, Dahomey Gap, western Central Africa, Gabon and Central Africa (Fig. 1). All the sequences produced from Togo, Benin and southwestern Nigeria clustered into the Dahomey Gap lineage (bootstrap support = 75%).

**Figure 1.**
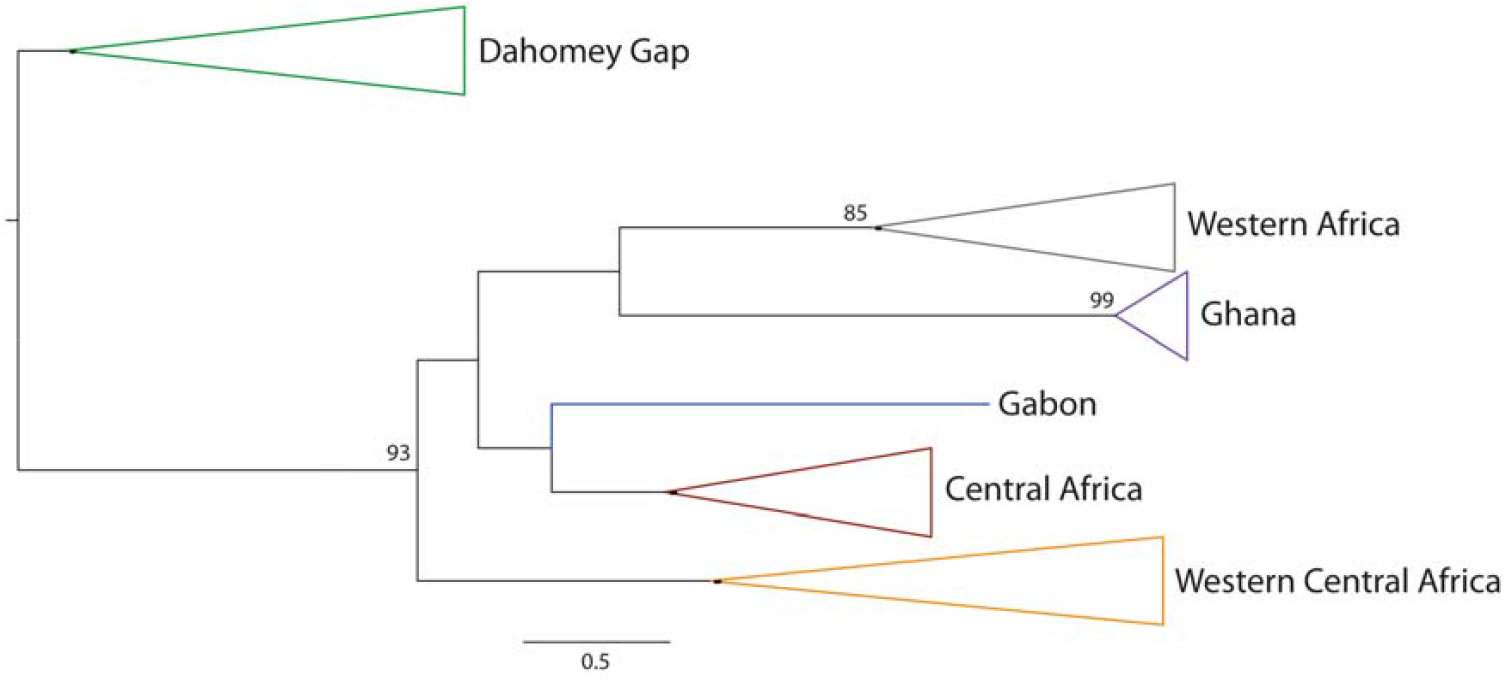
Neighbor-joining tree of white-bellied pangolins based on 268 control region sequences, showing the six main geographic lineages (following Gaubert et al. 2016) collapsed. Bootstrap supports are given at nodes. All the individuals from the Dahomey Gap belong to the Dahomey Gap lineage (see Fig S8, Supplementary information, for the expanded tree).

We identified fourteen CR1 haplotypes in DGL. The median-joining network did not show any specific geographic structure in haplotype distribution (Fig S1, Supplementary information). Two haplotypes were dominant in the DGL, with H5 (45%) being widely distributed and H1 (25%) located in the central and northern part of the range (Fig. 2). Nine haplotypes were found both in forests and wildlife markets, while four were only found in wildlife markets. The proportion of H5 (40 %) and H1 (27%) in wildlife markets was reflective of their frequencies observed in forest populations.

**Figure 2.**
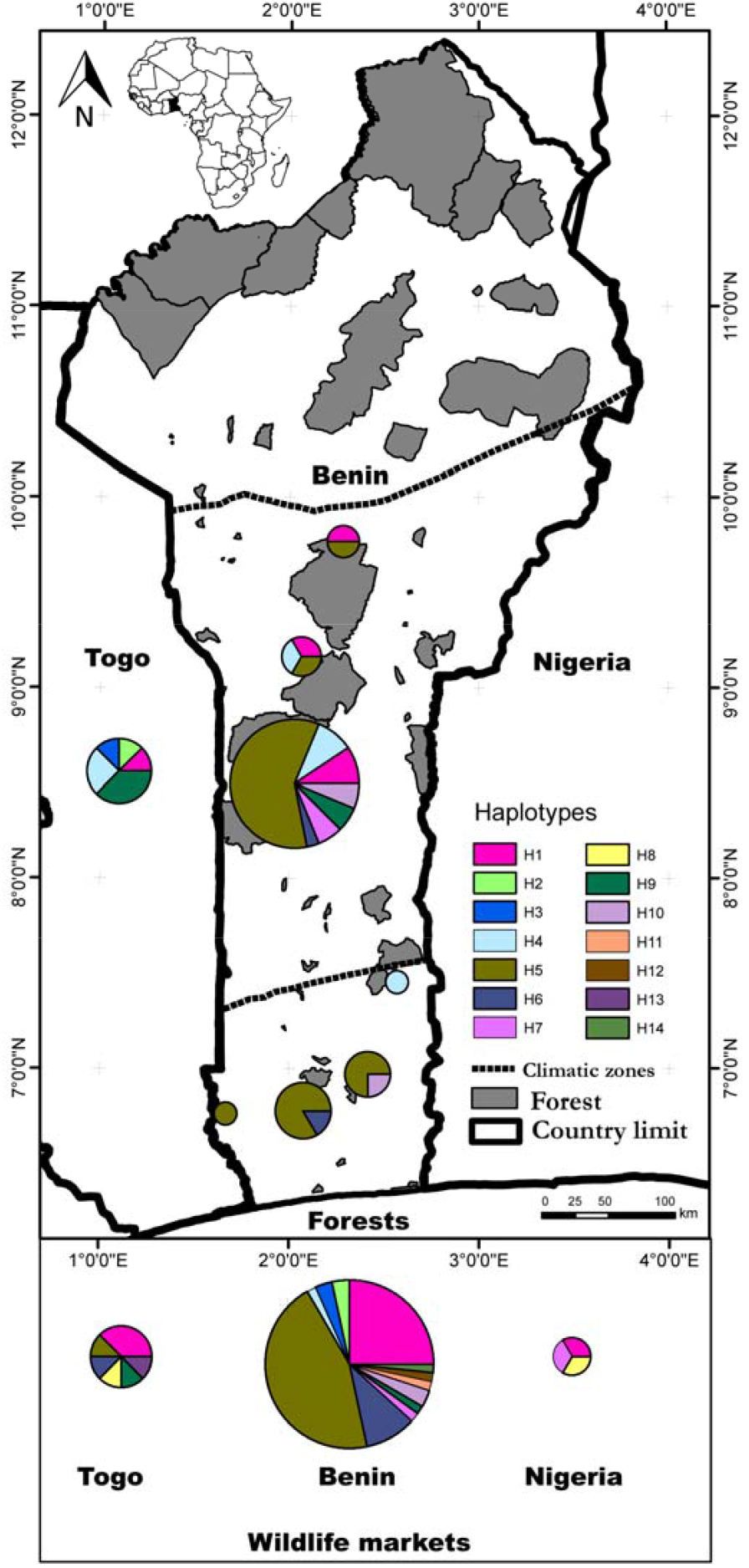
Distribution of control region haplotypes in white-bellied pangolins from the forests and wildlife markets of the Dahomey Gap. Haplotype numbers refer to Table S2 (Supplementary information). Top left shows the location of the study zone in Africa (in black).

Mismatch analysis of CR1 haplotypes showed a bimodal distribution significantly deviating from the sudden expansion demographic model (P(Sim. Rag. >= Obs. Rag.) = 0.005), whereas the spatial expansion model could not be rejected (P(Sim. Rag. >= Obs. Rag.) = 0.15). The statistics D (-0.56496), *r* (0.2057) and R2 (0.0710) showed no significant deviation from a scenario of large and constant population size through time (*p >* 0.10), whereas Fs (-4.780, *p* = 0.0431) significantly rejected the model.

### Microsatellites

Within DGL, the number of alleles (Na) varied from 2 to 11 (mean=5.3). Allelic richness (A_R_) ranged from 1.86 to 8.04 (mean = 4.27; sample size = 169), and observed heterozygosity (*Ho*) and expected heterozygosity (*He*) from 0.072 to 0.775 (mean *=* 0.414) and 0.069 to 0.842 (mean *=* 0.498), respectively. Eleven loci deviated significantly from *HWE* (*P*<0.05). Six of them showed significant levels of heterozygosity deficiency (*P <* 0.0025) and three of them were involved in LD (PT_839522, PT_1453906 and PT_353755). Null alleles were identified in ten loci, including seven that deviated from *HWE* (Table 1).

**Table 1.** Genetic diversity estimates at the 20 microsatellite loci used in this study. *N* = number of genotyped individuals; *Na* = number of alleles; *A*_*R*_ = allelic richness; *H*_*O*_ = observed heterozygosity; *H*_*E*_ = expected heterozygosity; *P(HWE)* = *P*-value for deviation from the null hypothesis of Hardy–Weinberg equilibrum; *F*_*IS*_ = inbreeding coefficient. *P*-values *\* <0*.*05, *** <0*.*001*.

| Locus | $N$ | $Na$ | $A_R$ | $H_O$ | $H_E$ | $p(HWE)$ | $F_{IS}$ |
| --- | --- | --- | --- | --- | --- | --- | --- |
| **PT_1162028** | 169 | 4 | 3.123 | 0.426 | 0.6 | 0.000*** | 0.292* |
| **PT_1753627** | 162 | 11 | 7.993 | 0.642 | 0.689 | 0.019* | 0.071 |
| **PT_796077** | 168 | 3 | 2.117 | 0.16 | 0.186 | 0.281 | 0.145 |
| **PT_1973508** | 169 | 5 | 4.339 | 0.562 | 0.48 | 0.028* | -0.167 |
| **PT_839522** | 168 | 7 | 6.168 | 0.524 | 0.788 | 0.000*** | 0.338* |
| **PT_464918** | 168 | 8 | 7.025 | 0.56 | 0.766 | 0.000*** | 0.272* |
| **PT_1453906** | 168 | 4 | 3.851 | 0.321 | 0.646 | 0.000*** | 0.505* |
| **PT_34432** | 151 | 2 | 1.863 | 0.086 | 0.082 | 0.580 | -0.042 |
| **PT_1594892** | 151 | 3 | 2.988 | 0.192 | 0.501 | 0.000*** | 0.619* |
| **PT_308752** | 169 | 11 | 7.203 | 0.657 | 0.678 | 0.000*** | 0.034 |
| **PT_1669238** | 169 | 3 | 2.233 | 0.314 | 0.361 | 0.276 | 0.134 |
| **PT_1225378** | 167 | 3 | 1.903 | 0.072 | 0.069 | 0.972 | -0.031 |
| **PT_739516** | 167 | 4 | 3.363 | 0.461 | 0.498 | 0.612 | 0.077 |
| **PT_619913** | 168 | 2 | 1.959 | 0.113 | 0.128 | 0.142 | 0.116 |
| **PT_338821** | 165 | 7 | 5.930 | 0.503 | 0.721 | 0.000*** | 0.305* |
| **PT_18497228** | 168 | 4 | 3.848 | 0.595 | 0.6 | 0.611 | 0.01 |
| **PT_378852** | 168 | 2 | 1.969 | 0.113 | 0.138 | 0.020* | 0.182 |
| **PT_353755** | 160 | 9 | 8.044 | 0.775 | 0.842 | 0.000* | 0.083 |
| **PT_276641** | 166 | 5 | 4.168 | 0.699 | 0.696 | 0.535 | -0.002 |
| **PT_2019332** | 169 | 8 | 5.319 | 0.503 | 0.498 | 0.309 | -0.007 |

Genetic variance within DGL individuals did not show any geographic structuring on the main PCoA axes (PC1 to PC3 = 12.36 % of total variance), whether individuals from wildlife markets were considered or not (Fig. 3).

**Figure 3.**
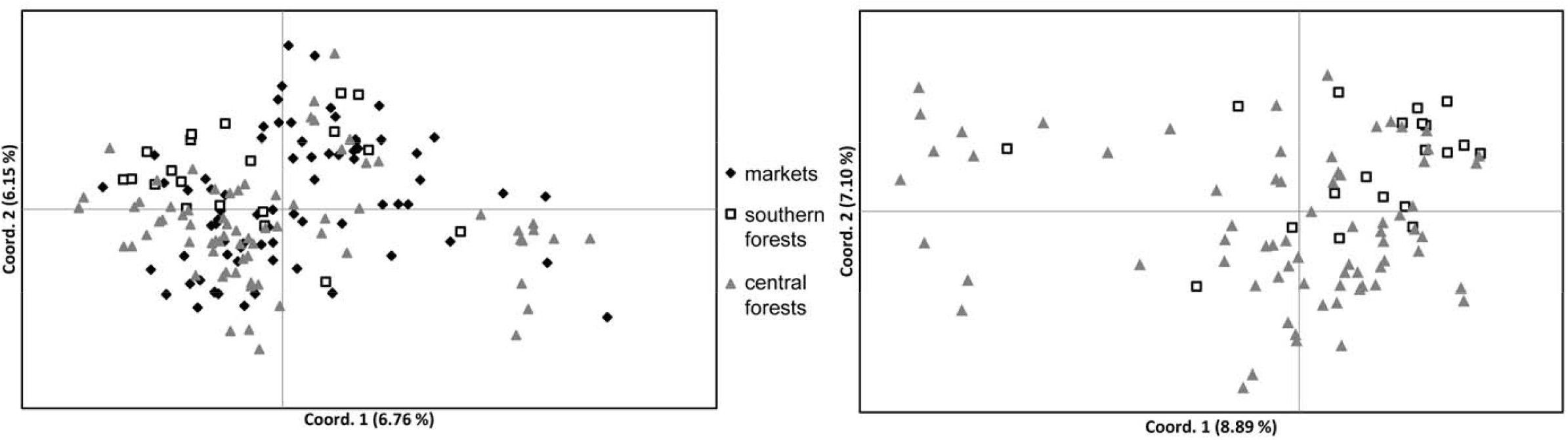
Distribution of genetic variance (PCoA) within white-bellied pangolins from the Dahomey Gap lineage. A- including forests and wildlife markets; B- excluding wildlife markets.

Paiwise population differentiations using partition schemes (ii) and (iii) ranged from low (*F*_*ST*_ = 0.0378) to moderate (*F*_*ST*_ = 0.116; between central Togo and southern Benin). In partition scheme (i), differentiations among the six forest populations were the greatest and all significant (*p*<0.05) and ranged from low to moderate (*F*_*ST*_ = 0.0528-0.1399; 67% of the *F*_*ST*_ values) and high (0.166 < *F*_*ST*_ < 0.244; 27% of the *F*_*ST*_ values) (Table S3, Supplementary information). The mean inbreeding coefficient (*F*_*IS*_) in the six DGL populations was 0.172 and varied from 0.098 to 0.317 (Table S4, Supplementary information).

Bayesian clustering analysis with STRUCTURE –without prior information on locations– detected a most likely number of populations K = 3 (Fig S2, Supplementary information). The three clusters did not correspond to exclusive geographic delineations, each being an admixture of individuals from southern and central forest regions together with wildlife markets. The assignment probabilities of the individuals to their respective populations were generally low. Under the LOCPRIOR model, r was equal to 2.368.

Final inference of population number using *Geneland* was K = 7 (95% of the runs). The geographic delineation of the populations did not show any spatially exclusive distribution. Assignment probabilities to populations varied greatly among the best five runs (data not shown). Similar results were observed when fixing K = 3 (Fig S4, Supplementary information).

The eigenvalues observed from the spCA analysis (Fig. 4a) suggested a relatively strong signal of “local structure”, indicating negative autocorrelation between geographic and genetic distances in pangolins from the Dahomey Gap. However, we could not detect any significant global or local structure signal (*p* = 0.602 and 0.102, respectively) across the study area.

**Figure 4.**
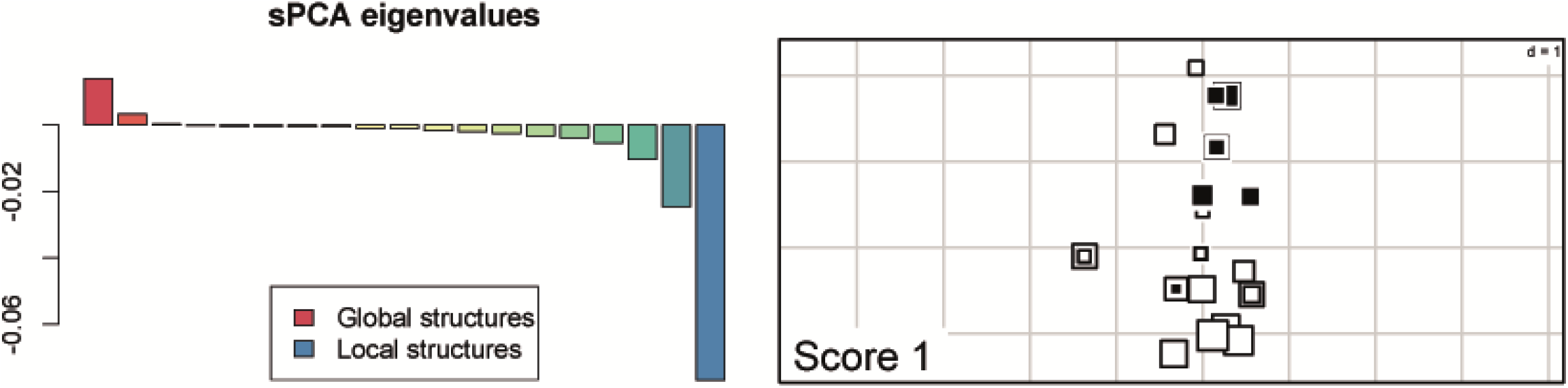
Spatial clustering in white-bellied pangolins from the Dahomey Gap obtained from spatial Principal Component Analysis (sPCA). Left: Positive and negative sPCA eigenvalues indicating global and local structures, respectively. Right: Map of the first global sPCA score among sampling localities. Large white and black squares stand for highly negative and positive scores respectively. Large white squares are genetically well differentiated from large black squares, while small squares are less differentiated.

There was significant IBD effect among forest individuals (*r* = 0.128; *p* = 0.001) and between populations (forests with at least 7 individuals) (*r* = 0.779; *p* = 0.009) (Fig. 5).

**Figure 5.**
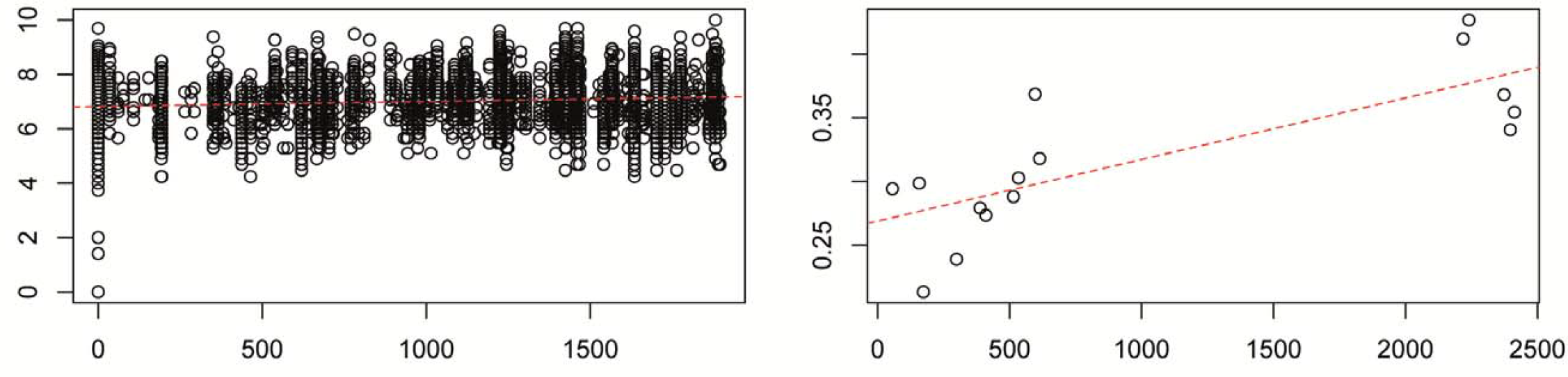
Isolation by distance among (left) individuals and (right) populations of white-bellied pangolins from the Dahomey Gap as inferred from 20 microsatellite loci. Dashed curve indicates linear regression.

A total of 164 samples (97 %) had a unique genotype. Five samples shared a genotype with four other samples, including three pairs (A65-A66, B1-B12, B2-B15) and one triplet (D14-D15-D16; see Table S1, Supplementary information). The null hypothesis of encoutering the different genotypes more than once by chance was rejected in all cases (P < 0.0001). The unbiased probability of identity (uPI) and the probability of identity among siblings (PIsibs) were both low (uPI= 8.12 e^-13^; PIsibs = 9.22 e^-06^). A number of 7 microsatellite loci were needed to reach the conservative value of PIsibs < 0.01 (Fig S5, Supplementary information).

Estimated mean frequencies of private alleles across the 20 loci (sample size = 7) using ADZE ranged from 0.10 to 0.24. Within the eleven loci that presented appropriate private allele signatures for one or several populations (see Fig S6 and S7, Supplementary information), seven loci provided six observed private alleles in GenAlEx that could potentially differentiate among four populations. On this basis, five individuals found on wildlife markets could be traced back to their forests of origin: A2 (Gbèdagba market, Central Benin) and A38 (Azovè market, southern Benin) to the Wari-Maro forest reserve, B50 (Azovè market, southern Benin) to the forests of central Togo, and D13 (Avogbannan market) and D26 (Dantokpa market) to the Ouémé supérieur forest reserve.

Bottleneck analysis on the DGL across 20 loci was not significant (Wilcoxon sign-rank test; *p*>0.05) for both SMM and TPM models.

*VarEff* identified a pronounced and recent decline in the effective population size (*Ne*) of DGL regardless of the models (Fig. 6). Our results suggested a 92-98% reduction of *Ne*, from 1682-3440 (ancestral *Ne*) to 78-135 (contemporaneous *Ne*) individuals as harmonic means (95% CI: 263 to 487). The decrease in *Ne* was estimated to occur c. 200-500 years ago.

**Figure 6.**
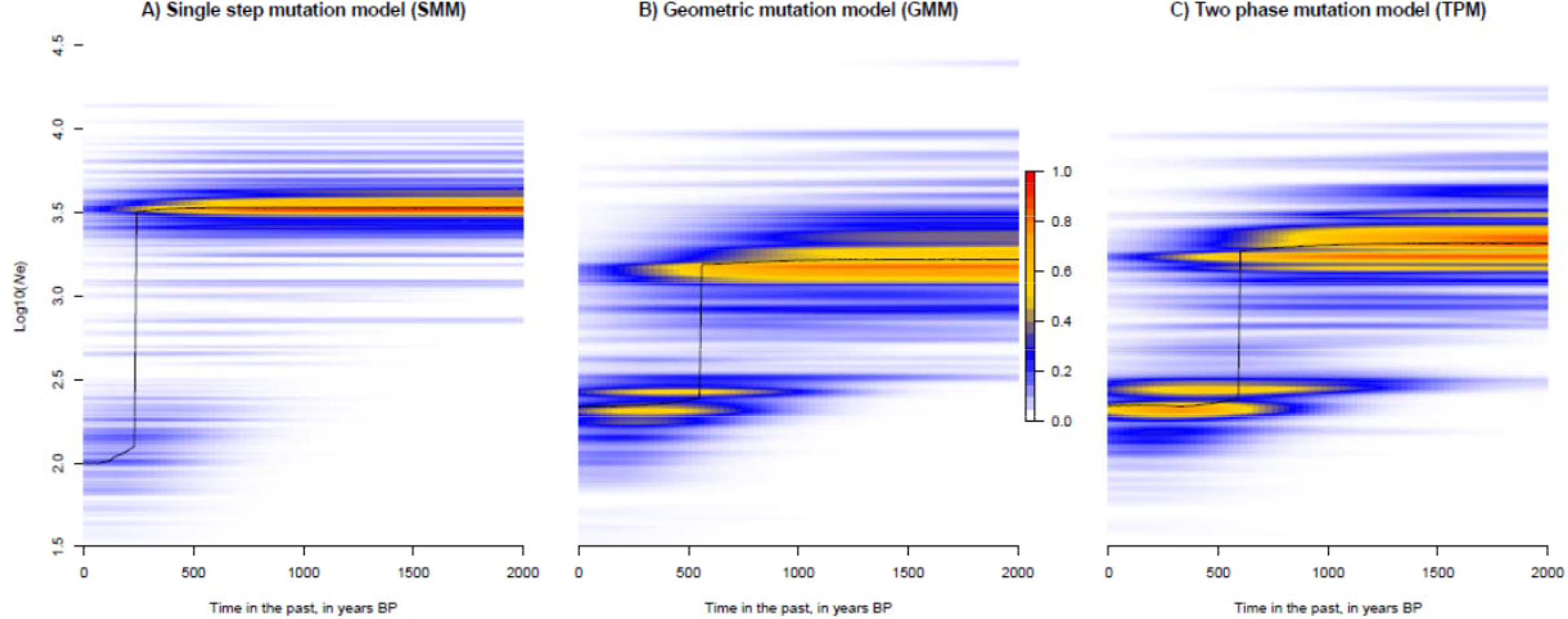
Temporal change in the effective population size of white-bellied pangolins in the Dahomey Gap, as estimated from *VarEff* under three different mutation models. Mode (black line) and kernel density (color scale) of effective population size (*Ne*) posterior distributions are given in years BP.

## Discussion

### The endemic lineage of white-bellied pangolins from the Dahomey Gap feeds an endemic trade

Our mtDNA tree-based assignment procedure showed that all the white-bellied pangolins (WBP) collected and traded in the Dahomey Gap –from Togo, Benin and southwestern Nigeria– belong to the Dahomey Gap lineage (DGL; [9]). With 168 new samples sequenced from different forests and TMM, we have circumbscribed with more accuracy the geographic delimitation of the DGL, from central Togo to northernmost and southernmost locations in Benin (Ouémé supérieur and Gnanhouizounmè, respectively), and Asejire in southwestern Nigeria. We have also confirmed the absence of range overlap with other WBP lineages, notably from the neighboring Ghana. This last result is to be tempered by the fact that introgression between WBP lineages cannot be excluded –but has not been reported to date– and could have passed undetected for the few samples sequenced at a single locus (mtDNA) in our study. The endemic pattern of DGL superimposes with DNA-based delineation recently found in mammals and plants from the Dahomey Gap [52, 53], further emphasizing the patrimonial importance of the area for West African forest taxa.

Contrary to the bushmeat markets in Têgon and Hounkpogon (Benin) and Asejire (southwestern Nigeria) that are known to source the game from nearby forests [18], this study), the endemicity of the pangolin trade was not expected from the traditional medicine markets (TMMs). Indeed, the geographic amplitude of the TTM network was shown in previous investigations from Benin and Nigeria, revealing the long-distance trade of non-native species [54, 55]. Moreover, there is a great demand for pangolins in the Dahomey Gap [56], notably from the Chinese diaspora [7], and the trade of pangolins across borders have been reported elsewhere in tropical Africa [57]. The endemicity of the pangolin trade in the Dahomey Gap might translate into a huge hunting pressure on such a geographically restricted lineage. This is especially true as what we observed on the markets might not apprehend the full dimension of the pangolin trade, which may also feed the international market through alternative networks [7, 57, 58].

### White-bellied pangolins in the Dahomey Gap show genetic diversity erosion and recent, sharp demographic decline

Overall, DGL populations were characterized by low levels of genetic diversity. Mitochondrial (CR1) haplotype and nucleotide diversity was lower compared to all the other WBP lineages. Mean allelic richness based on microsatellites was also lower than what was found in WBP from Cameroon using the same markers (*A*_*R*_ = 4.63 *vs*. 6.74, respectively; minimum sample size = 37; see [17]). Compared to genetic diversity estimates based on ten WBP samples from Ghana, mean observed heterozygosity was again lower in DGL (*Ho* = 0.541 *vs*. 0.414, respectively; see [59]).

Low levels of genetic diversity are assumed to be negatively correlated with fitness and adaptability [60]. The overall absence of equilibrium detected from our microsatellite dataset suggests that inbreeding depression is one of the driving factors of the low genetic diversity observed in DGL. Although deviations from Hardy-Weinberg equilibrium (55% of the loci in this study) can be due to a number of factors including inbreeding, population structure and genotyping errors [61], we can reasonably discard the two latter given (i) the apparent lack of population structure in DGL and (ii) the optimized loci and genotyping approach that we used. Besides, it is well known that populations going through inbreeding will produce upwardly biased estimates of null allele frequencies [62], as observed in DGL (50% of the loci). The deficit of heterozygotes observed in 75% of the loci, 30% of which having significant levels of deficiency, also supports the view that DGL populations are subject to inbreeding depression, possibly indicative of non-random mating [63].

Our demographic analyses based on microsatellites identified a sharp and recent decline in the effective population size (*Ne*) of DGL c. 200-500 years ago (100-250 generations), leading to a 92-98 % reduction to contemporaneous *Ne* below the conservative thresholds of minimum viable population size (500-5000; [64, 65]). As variation in *N*_*e*_ is crucial in determining levels of genetic diversity [66], the state of genetic depauperation observed in DGL may be directly linked to the recent demographic decline affecting the lineage. The time of decline corresponds to a period of major transformations along the ‘Slave Coast’, where from the 17^th^ century the Dahomey kingdom expanded as a state bureaucracy benefiting from the growing trade of slaves and agricultural goods with Europeans [67, 68]. Whether such political growth was followed by agricultural expansion and deforestation causing the decline of pangolins in the region is uncertain, but similar declines have been observed in commercially exploited species of vertebrates through the last centuries [68–70]. Our results are important for the conservation of DGL, because inbreeding depression together with high levels of genetic drift will potentially lead to the fixation of mildly deleterious alleles that could drive an extinction vortex in this lineage [71].

Earlier events such as the expansion of agriculture in West Africa c. 4,200 BP [72] and natural forest fragmentation caused by cyclical drier climatic conditions in the Dahomey Gap from 4,500 BP [10] do not seem to have affected the demographic history of DGL populations. Indeed, mtDNA-based demographic analyses showed no deviation from a model of large, constant *Ne* through time, indicating long-term matrilineal stability. The only exception was the Fu’s statistics, which has maximum power to detect sudden demographic decline events [73] and thus can be related to the recent decline discussed above. However, further analyses based on nuclear genomic markers (SNPs) will have to be conducted to assess the ancient demographic history of DGL, the origin of which dates back to 120-240 kya [9]. The Dahomey Gap is a broad savannah corridor intermixed with forest patches that seperates the two African rainforest blocks, and as such can be considered a sub-optimal habitat for WBP that heavily relies on rainforest cover [9]. Because the area underwent drastic alternations of dry and humid periods since at least the last 150,000 yr [74], it is possible that DGL populations were affected by early, successive founder effects and bottlenecks due to Late Pleistocene climatic pejoration [53]. Such demographic events could also have shaped the genetic diversity and absence of population structure (see below) observed today in DGL.

### The fragmented populations of white-bellied pangolins in the Dahomey Gap show no genetic structure

Our analyses based on mtDNA and microsatellites generally suggested that there was no geographic structuring across the Dahomey Gap, against the expectation that habitat fragmentation leads to genetic isolation [75]. One potential explanation to the lack of population structure would involve a fair level of long-range dispersals. Pangolins have been reported to disperse up to 300 km in four months, with a marked period of mobility for unestablished young individuals through –notably– anthropized areas [76, 77] However, the dispersal ecology of pangolins remains poorly known, especially in WBP. The species seems to heavily rely on forest cover and old trees for its nocturnal activities [78], exploring its range up to 1.8 km per night [79]. In the Dahomey Gap, WBP may occur in disturbed habitats including commercial plantations of teaks and palm trees, fallows and farmlands [80]. However, evidence of long-range dispersal is lacking and the general absence of structural connectivity among the remnant forest islands of the Dahomey Gap, especially in Benin (see [81, 82]) does not support such scenario.

Although we could not find any clear population structure in DGL, we detected significant levels of differentiation and isolation-by-distance among both individuals and populations. We observed the strongest differentiations between the most distant Togolese and Beninese populations, and some cases of moderate differentiation between geographically close populations (e.g., the contiguous Ouémé supérieur and Wari-Maro protected areas). Genetic differentiation among populations is determined by the interplay between homogenizing processes such as gene flow and differentiating processes including local adaptation, different adaptive responses to shared environmental conditions, and genetic drift [83]. If we posit that, in the case of DGL, (i) gene flow between populations is not an option (see above), (ii) genetic drift in isolated, inbred populations should have resulted in detectable geographic structure [84], and (iii) environmental conditions are similar across the DGL range (Guineo-Congolian and Sudano-Guinean zones; [85]), then different adaptive responses of populations to a similar environment could be a candidate scenario to explain the level of differentiation observed among populations from the Dahomey Gap. This could notably be expectable in the case of populations trapped into sub-optimal environmental conditions and impacted by frequent climatic oscillations (see [9]). However, microsatellites markers generally reflect neutral genetic variation, and should not be affected by signatures of local adaptation [86].

A more plausible scenario would relate to a former, possibly recent, spatial expansion in subdivided populations across the DGL range, as mtDNA did not reject this model. It is possible that forest-restricted DGL populations underwent a spatial expansion following the last recent increase in rainforest cover during the last Interglacial or early Holocene periods [10]. Such event would explain the absence of population structure together with some level of population differentiation as observed in this study, provided that dispersal among populations would have been maintained long enough to counter-balance the effect of genetic drift in isolated populations later induced by drier climatic conditions,[10]) and anthropogenic pressure in the Dahomey Gap. Overall, our study has the merit to posit a number of hypotheses that could explain the puzzling pattern observed in the population structure of DGL. Such hypotheses will have to be tested through a demographic scenario-based strategy, preferentially using versatile and powerful genomic resources.

### Tracing the pangolin trade in the Dahomey Gap: specimen dispatch and evidence for long-distance trade

The genetic diversity of WBP sold on the markets was reflective of the overall genetic diversity (haplotype and allelic frequencies) observed in DGL forest populations, suggesting a widespread sourcing of pangolins through the entire Dahomey Gap. Our 20 microsatellites loci provided the necessary power to confidently distinguish among all the DGL indivuals, and only seven best microsatellite loci were needed to reach the conservative value of probability of identity < 0.01 [87]. The probability that two individuals drawn at random from a population, including or not including siblings, will have the same genotype was low (but higher than in previous studies,[17, 77]). This has important implications for the genetic tracing of the pangolin trade in the Dahomey Gap, as one of the main inputs of the genetic toolkit is its potential for tracing the trade at the individual level [88]. For instance, our markers would be capable of estimating the exact number of individuals from scale seizures, a major challenge that conservationists are currently facing [89]. We also demonstrated that our genotyping approach was useful in tracing the dispatch of a same pangolin sold on the market (one individual detected on two different stalls in Dantokpa) or kept by local communities (scales of two different individuals shared between villagers in Mont Kouffé and Lama forests). Such application is especially relevant for tracing the pangolin trade, which often translates into separate networks specialized in the selling of specific parts (scales, organs, meat), notably in Benin [7].

Given the lack of population structure in DGL and the negative autocorrelation between geographic and genetic distances (for individuals and populations), classical assignment procedures [90, 91] are hardly usable to trace the pangolin trade in the Dahomey Gap. We have developed a conservative approach combining rarefaction analysis of private allele frequencies in each population and cross-validation with observed data that could partly circumvent that issue. Such method may be applicable to any taxon or lineage without observed genetic structure across its range, notably at the local scale. We identified seven private alleles (from seven loci) that could potentially differentiate among five DGL populations. Five pangolins were traced to their forest of origin using three private alleles, illustrating the long-distance trading routes that feed TMMs in the Dahomey Gap (see [7]). Indeed, pangolins from the forests of central Togo, Wari-Maro and Ouémé Supérieur were found on the markets of Dantokpa, Gbèdagba, Azovè and Avogbannan, c. 200-300 km away from the source forests.

Despite a relatively fair number of loci (20) and an exhaustive sample set across Benin, we could only trace c. 8% of the WBP genotyped from TMMs. Such performance could be improved with additional geographic sampling, notably from Togo and southwestern Nigeria, and denser sampling of populations, although the ideal standards for reaching confident estimates of allele frequencies when applied to forensic use might be unreachable (100-150 individuals per population; [92]). Future analyses based on bi-allelic markers such as SNPs will have to be considered as they significantly reduce the sample size required for reliable estimates of allele frequency distribution [93].

## Conclusion

Our study is the first to provide a comprehensive population genetic assessment of an African pangolin species/lineage, filling an important knowledge gap for the future conservation of pangolins in Africa [16]. Overall, we showed that the DGL populations suffered from inbreeding, genetic diversity erosion and a drastic decline in effective population size. Given the multi-purpose trade that DGL populations are the target of (this study, [7]), and the observed reduction of the DGL range during the last two decades (in Benin, [11]), the conservation status of WBP in the Dahomey Gap should be urgently re-assessed. Conservation measures are to be implemented before the species becomes locally extinct, as it may already be the case for the giant pangolin [11]. Measures should include the reinforcement of national protection status, the creation of dedicated protected areas and forest corridors, campaigns of public awareness, and breeding programs.

## Supporting information

Supplemental list

## Acknowledgments

We are grateful to the Direction Générale des Eaux-Forêts et Chasse of Benin for providing the research permits to conduct fieldwork. We are grateful to the staff of the Plateau technique - Biologie moléculaire et microbiologie at EDB for assistance during lab work. We thank all the local hunters and vendors in the TMMs who provided us with the necessary tissue samples by their free consent.

## References

1. Challender DWS, Heinrich S, Shepherd CR, Katsis LKD. Chapter 16 - International trade and trafficking in pangolins, 1900–2019. In: Challender DWS, Nash HC, Waterman C, editors. Pangolins. Academic Press; 2020. p. 259–76. doi:10.1016/B978-0-12-815507-3.00016-2.

2. Frutos R, Serra-Cobo J, Chen T, Devaux CA. COVID-19: Time to exonerate the pangolin from the transmission of SARS-CoV-2 to humans. Infect Genet Evol. 2020;84:104493.

3. Aditya V, Goswami R, Mendis A, Roopa R. Scale of the issue: Mapping the impact of the COVID-19 lockdown on pangolin trade across India. Biol Conserv. 2021;257:109136.

4. Ingram DJ, Cronin DT, Challender DWS, Venditti DM, Gonder MK. Characterising trafficking and trade of pangolins in the Gulf of Guinea. Glob Ecol Conserv. 2019;17:e00576.

5. Pietersen D, Moumbolou C, Ingram DJ, Soewu D, Jansen R, Sodeinde D, et al. IUCN Red List of Threatened Species: Phataginus tricuspis. IUCN Red List Threat Species. 2019. https://www.iucnredlist.org/en. Accessed 8 Aug 2021.

6. Mambeya MM, Baker F, Momboua BR, Pambo AFK, Hega M, Okouyi VJO, et al. The emergence of a commercial trade in pangolins from Gabon. Afr J Ecol. 2018;56:601–9.

7. Zanvo S, Djagoun SCAM, Azihou FA, Djossa B, Sinsin B, Gaubert P. Ethnozoological and commercial drivers of the pangolin trade in Benin. J Ethnobiol Ethnomed. 2021;17:18.

8. Zhang H, Ades G, Miller MP, Yang F, Lai K, Fischer GA. Genetic identification of African pangolins and their origin in illegal trade. Glob Ecol Conserv. 2020;23:e01119.

9. Gaubert P, Njiokou F, Ngua G, Afiademanyo K, Dufour S, Malekani J, et al. Phylogeography of the heavily poached African common pangolin (Pholidota, Manis tricuspis) reveals six cryptic lineages as traceable signatures of Pleistocene diversification. Mol Ecol. 2016;25:5975–93.

10. Salzmann U, Hoelzmann P. The Dahomey Gap: an abrupt climatically induced rain forest fragmentation in West Africa during the late Holocene. The Holocene. 2005;15:190–9.

11. Zanvo S, Gaubert P, Djagoun CAMS, Azihou AF, Djossa B, Sinsin B. Assessing the spatiotemporal dynamics of endangered mammals through local ecological knowledge combined with direct evidence: The case of pangolins in Benin (West Africa). Glob Ecol Conserv. 2020;23:e01085.

12. Segniagbeto GH, Assou D, Agbessi EKG, Atsri HK, D’Cruze N, Auliya M, et al. Insights into the status and distribution of pangolins in Togo (West Africa). Afr J Ecol. 2021;59:342–9.

13. Omifolaji JK, Ikyaagba ET, Jimoh SO, Ibrahim AS, Ahmad S, Luan X. The emergence of Nigeria as a staging ground in the illegal pangolin exportation to South East Asia. Forensic Sci Int Rep. 2020;2:100138.

14. Lorenzini R, Fanelli R, Tancredi F, Siclari A, Garofalo L. Matching STR and SNP genotyping to discriminate between wild boar, domestic pigs and their recent hybrids for forensic purposes. Sci Rep. 2020;10:3188.

15. Lorenzini R, Garofalo L. Wildlife forensics: DNA analysis in wildlife forensic investigations. In: Forensic DNA Analysis. Apple Academic Press; 2021.

16. Heighton SP, Gaubert P. A timely systematic review on pangolin research, commercialization, and popularization to identify knowledge gaps and produce conservation guidelines. Biol Conserv. 2021;256:109042.

17. Aguillon S, Din Dipita A, Lecompte E, Missoup AD, Tindo M, Gaubert P. Development and characterization of 20 polymorphic microsatellite markers for the white-bellied pangolin Phataginus tricuspis (Mammalia, Pholidota). Mol Biol Rep. 2020;47:1–7.

18. Olayemi A, Oyeyiola A, Antunes A, Bonillo C, Cruand C, Gaubert P. Contribution of DNA-typing to bushmeat surveys: Assessment of a roadside market in south-western Nigeria. Wildl Res. 2011;38:696.

19. Gaubert P, Zenatello M. Ancient DNA perspective on the failed introduction of mongooses in Italy during the XXth century. J Zool. 2009;279:262–9.

20. Hall TA. Bioedit: a user friendly biological sequence alignment editor and analysis pogramme for Windows 95/98/NT. Nucleic Acids Symp Serv. 1999;41:95–8.

21. Taberlet P, Griffin S, Goossens B, Questiau S, Manceau V, Escaravage N, et al. Reliable genotyping of samples with very low DNA quantities using PCR. Nucleic Acids Res. 1996;24:3189–94.

22. Kumar S, Stecher G, Li M, Knyaz C, Tamura K. MEGA X: Molecular evolutionary genetics analysis across computing platforms. Mol Biol Evol. 2018;35:1547–9.

23. Kimura M. A simple method for estimating evolutionary rates of base substitutions through comparative studies of nucleotide sequences. J Mol Evol. 1980;16:111–20.

24. Rozas J, Ferrer-Mata A, Sánchez-DelBarrio JC, Guirao-Rico S, Librado P, Ramos-Onsins SE, et al. DnaSP 6: DNA sequence polymorphism analysis of large data sets. Mol Biol Evol. 2017;34:3299–302.

25. Excoffier L, Lischer HEL. An integrated software package for population genetics data analysis. 2015. http://cmpg.unibe.ch/software/arlequin35/. Accessed 13 Aug 2021.

26. Schneider S, Excoffier L. Estimation of past demographic parameters from the distribution of pairwise differences when the mutation rates vary among sites: Application to human mitochondrial DNA. Genetics. 1999;152:1079–89.

27. Tajima F. Statistical method for testing the neutral mutation hypothesis by DNA polymorphism. Genetics. 1989;123:585–95.

28. Fu YX. Statistical tests of neutrality of mutations against population growth, hitchhiking and background selection. Genetics. 1997;147:915–25.

29. Harpending HC. Signature of ancient population growth in a low-resolution mitochondrial DNA mismatch distribution. Hum Biol. 1994;66:591–600.

30. Ramos-Onsins SE, Rozas J. Statistical properties of new neutrality tests against population growth. Mol Biol Evol. 2002;19:2092–100.

31. Hudson RR. Gene genealogies and the coalescent process. Oxf Surv Evol Biol. 1990;7:1–44.

32. Kearse M, Moir R, Wilson A, Stones-Havas S, Cheung M, Sturrock S, et al. Geneious Basic: An integrated and extendable desktop software platform for the organization and analysis of sequence data. Bioinformatics. 2012;28:1647–9.

33. Peakall R, Smouse PE. GenAlEx 6.5: genetic analysis in Excel. Population genetic software for teaching and research—an update. Bioinformatics. 2012;28:2537–9.

34. Goudet J. FSTAT (version 2.9. 4), a program (for Windows 95 and above) to estimate and test population genetics parameters. http://www2.unil.ch/popgen/softwares/fstat.htm. 2003.

35. Oosterhout CV, Hutchinson WF, Wills DPM, Shipley P. micro-checker: software for identifying and correcting genotyping errors in microsatellite data. Mol Ecol Notes. 2004;4:535–8.

36. Excoffier L, Lischer HEL. Arlequin suite ver 3.5: a new series of programs to perform population genetics analyses under Linux and Windows. Mol Ecol Resour. 2010;10:564–7.

37. Pritchard JK, Wen X, Falush D. Documentation for structure software: Version 2.3. 2010;39.

38. Earl DA, vonHoldt BM. STRUCTURE HARVESTER: A website and program for visualizing STRUCTURE output and implementing the Evanno method. Conserv Genet Resour. 2012;4:359–61.

39. Evanno G, Regnaut S, Goudet J. Detecting the number of clusters of individuals using the software structure: a simulation study. Mol Ecol. 2005;14:2611–20.

40. Guillot G, Mortier F, Estoup A. Geneland: a computer package for landscape genetics. Mol Ecol Notes. 2005;5:712–5.

41. Jombart T. adegenet: a R package for the multivariate analysis of genetic markers. Bioinforma Oxf Engl. 2008;24:1403–5.

42. Jombart T. adegenet: a R package for the multivariate analysis of genetic markers. Bioinforma Oxf Engl. 2008;24:1403–5.

43. Paradis E. pegas: an R package for population genetics with an integrated-modular approach. Bioinforma Oxf Engl. 2010;26:419–20.

44. Kamvar ZN, Tabima JF, Grünwald NJ. Poppr: an R package for genetic analysis of populations with clonal, partially clonal, and/or sexual reproduction. PeerJ. 2014;2:e281.

45. Valière N. gimlet: a computer program for analysing genetic individual identification data. Mol Ecol Notes. 2002;2:377–9.

46. Szpiech ZA, Jakobsson M, Rosenberg NA. ADZE: a rarefaction approach for counting alleles private to combinations of populations. Bioinforma Oxf Engl. 2008;24:2498–504.

47. Piry S, Luikart G, Cornuet J-M. Computer note. BOTTLENECK: a computer program for detecting recent reductions in the effective size using allele frequency data. J Hered. 1999;90:502–3.

48. Nikolic N, Chevalet C. VarEff. Variation of effective size. Software VAREFF (package R in file.zip) and the documentation. 2014. https://archimer.ifremer.fr/doc/00177/28781/. Accessed 8 Aug 2021.

49. Zhang H, Miller MP, Yang F, Chan HK, Gaubert P, Ades G, et al. Molecular tracing of confiscated pangolin scales for conservation and illegal trade monitoring in Southeast Asia. Glob Ecol Conserv. 2015;4:414–22.

50. Zhang F, Wu S, Zou C, Wang Q, Li S, Sun R. A note on captive breeding and reproductive parameters of the Chinese pangolin, Manis pentadactyla Linnaeus, 1758. ZooKeys. 2016;618:129–44.

51. Schlötterer C. Evolutionary dynamics of microsatellite DNA. Chromosoma. 2000;109:365–71.

52. Colyn M, Hulselmans J, Sonet G, Oudé P, De Winter J, Natta A, et al. Discovery of a new duiker species (Bovidae: Cephalophinae) from the Dahomey Gap, West Africa. Zootaxa. 2010;:1–30.

53. Demenou BB, Piñeiro R, Hardy OJ. Origin and history of the Dahomey Gap separating West and Central African rain forests: insights from the phylogeography of the legume tree Distemonanthus benthamianus. J Biogeogr. 2016;43:1020–31.

54. Saidu Y, Buij R. Traditional medicine trade in vulture parts in northern Nigeria. Vulture News. 2013;65:4–14.

55. Djagoun CAMS, Akpona H, Mensah G, Sinsin B, Nuttman C. Wild mammals trade for zootherapeutic and mythic purposes in Benin (West Africa): capitalizing species involved, provision sources, and implications for conservation. In: Animals in traditional folk medicine. 2013. p. 267–381.

56. D’Cruze N, Assou D, Coulthard E, Norrey J, Megson D, Macdonald DW, et al. Snake oil and pangolin scales: insights into wild animal use at “Marché des Fétiches” traditional medicine market, Togo. Nat Conserv. 2020;39:45–71.

57. Mambeya MM, Baker F, Momboua BR, Pambo AFK, Hega M, Okouyi VJO, et al. The emergence of a commercial trade in pangolins from Gabon. Afr J Ecol. 2018;56:601–9.

58. Ingram DJ, Coad L, Abernethy KA, Maisels F, Stokes EJ, Bobo KS, et al. Assessing Africa-wide pangolin exploitation by scaling local data. Conserv Lett. 2018;11:e12389.

59. du Toit Z, Dalton DL, du Plessis M, Jansen R, Grobler JP, Kotzé A. Isolation and characterization of 30 STRs in Temminck’s ground pangolin (Smutsia temminckii) and potential for cross amplification in other African species. J Genet. 2020;99:20.

60. Habel JC, Schmitt T. The burden of genetic diversity. Biol Conserv. 2012;147:270–4.

61. Wigginton JE, Cutler DJ, Abecasis GR. A Note on Exact Tests of Hardy-Weinberg Equilibrium. Am J Hum Genet. 2005;76:887–93.

62. Chybicki IJ, Burczyk J. Simultaneous estimation of null alleles and inbreeding coefficients. J Hered. 2009;100:106–13.

63. Keller LF, Waller DM. Inbreeding effects in wild populations. Trends Ecol Evol. 2002;17:230–41.

64. Reed DH, O’Grady JJ, Brook BW, Ballou JD, Frankham R. Estimates of minimum viable population sizes for vertebrates and factors influencing those estimates. Biol Conserv. 2003;113:23–34.

65. Clabby C. A Magic Number? Am Sci. 2010;98:24.

66. Charlesworth B. Effective population size and patterns of molecular evolution and variation. Nat Rev Genet. 2009;10:195–205.

67. Law R. Dahomey and the Slave Trade: Reflections on the historiography of the rise of Dahomey. J Afr Hist. 1986;27:237–67.

68. Monroe JC. The dynamics of state formation: The archaeology and ethnohistory of pre -colonial Dahomey - ProQuest. California; 2003. https://www.proquest.com/openview/5e43740457bdff3410ece1c79e26575f/1?pq-origsite=gscholar&cbl=18750&diss=y. Accessed 13 Aug 2021.

69. Bishop JM, Leslie AJ, Bourquin SL, O’Ryan C. Reduced effective population size in an overexploited population of the Nile crocodile (Crocodylus niloticus). Biol Conserv. 2009;142:2335–41.

70. Stoffel MA, Humble E, Paijmans AJ, Acevedo-Whitehouse K, Chilvers BL, Dickerson B, et al. Demographic histories and genetic diversity across pinnipeds are shaped by human exploitation, ecology and life-history. Nat Commun. 2018;9:4836.

71. Frankham R. Genetics and extinction. Biol Conserv. 2005;126:131–40.

72. Ozainne S, Lespez L, Garnier A, Ballouche A, Neumann K, Pays O, et al. A question of timing: spatio-temporal structure and mechanisms of early agriculture expansion in West Africa. J Archaeol Sci. 2014;50:359–68.

73. Ramírez-Soriano A, Ramos-Onsins SE, Rozas J, Calafell F, Navarro A. Statistical power analysis of neutrality tests under demographic expansions, contractions and bottlenecks with recombination. Genetics. 2008;179:555–67.

74. Dupont LM, Weinelt M. Vegetation history of the savanna corridor between the Guinean and the Congolian rain forest during the last 150,000 years. Veg Hist Archaeobotany. 1996;5:273–92.

75. Frankham R, Ballou JD, Ralls K, Eldridge MDB, Dudash MR, Fenster CB, et al. Genetic management of fragmented animal and plant populations. Oxford University Press; 2017. doi:10.1093/oso/9780198783398.001.0001.

76. Sun NC-M, Pei KJ-C, Lin J-S. Attaching tracking devices to pangolins: A comprehensive case study of Chinese pangolin Manis pentadactyla from southeastern Taiwan. Glob Ecol Conserv. 2019;20:e00700.

77. Ching-Min Sun N, Chang S-P, Lin J-S, Tseng Y-W, Jai-Chyi Pei K, Hung K-H. The genetic structure and mating system of a recovered Chinese pangolin population (Manis pentadactyla Linnaeus, 1758) as inferred by microsatellite markers. Glob Ecol Conserv. 2020;23:e01195.

78. Pages E. Les glandes odorantes des pangolins arboricoles (M. tricuspis et M. longicaudata): morphologie, developpement et roles. Biol Gabonica. 1968;4:353–400.

79. Pagès E. Eco-ethological study of Manis tricuspis by radio tracking. Mammalia. 1975;39:613–41.

80. Akpona HA, Djagoun CAMS, Sinsin B. Ecology and ethnozoology of the three-cusped pangolin Manis tricuspis (Mammalia, Pholidota) in the Lama forest reserve, Benin. Mammalia. 2008;72:198–202.

81. Mama A, Sinsin B, De Cannière C, Bogaert J. Anthropisation et dynamique des paysages en zone soudanienne au nord du Bénin. Tropicultura. 2013;31. https://orbi.uliege.be/handle/2268/160259. Accessed 14 Aug 2021.

82. Alohou EC, Gbemavo DSJC, Mensah S, Ouinsavi C. Fragmentation of forest ecosystems and connectivity between sacred groves and forest reserves in southeastern Benin, West Africa. Trop Conserv Sci. 2017;10:1940082917731730.

83. Whitlock MC. G’STand D do not replace FST. Mol Ecol. 2011;20:1083–91.

84. Lloyd MW, Campbell L, Neel MC. The power to detect recent fragmentation events using genetic differentiation methods. PLOS ONE. 2013;8:e63981.

85. White F. The vegetation of Africa: Descriptive memoir to accompany the UNESCO/AETFAT/UNSO vegetation map of Africa. Natural Resources Research Report XX. 7 Place de Fontenoy, 75700 Paris: UNESCO; 1983. https://www.creaf.uab.es/miramon/mmr/examples/miombo/docs/database/white/index.htm. Accessed 6 Oct 2021.

86. Holderegger R, Kamm U, Gugerli F. Adaptive vs. neutral genetic diversity: implications for landscape genetics. Landsc Ecol. 2006;21:797–807.

87. Waits LP, Luikart G, Taberlet P. Estimating the probability of identity among genotypes in natural populations: cautions and guidelines. Mol Ecol. 2001;10:249–56.

88. Nishant K, Vrijesh K, Ajay K. Wildlife forensic: current techniques and their limitations. J Forensic Sci Criminol. 2017;5:402.

89. Ullmann T, Veríssimo D, Challender DWS. Evaluating the application of scale frequency to estimate the size of pangolin scale seizures. Glob Ecol Conserv. 2019;20:e00776.

90. Hansen MM, Kenchington E, Nielsen EE. Assigning individual fish to populations using microsatellite DNA markers. Fish Fish. 2001;2:93–112.

91. Ogden R, Linacre A. Wildlife forensic science: A review of genetic geographic origin assignment. Forensic Sci Int Genet. 2015;18:152–9.

92. Chakraborty R. Sample size requirements for addressing the population genetic issues of forensic use of DNA typing. Hum Biol. 1992;64:141–59.

93. B-Rao C. Sample size considerations in genetic polymorphism studies. Hum Hered. 2001;52:191–200.

