## Supplemental list for "Can DNA help trace the local trade of pangolins? A genetic assessment of white-bellied pangolins from the Dahomey Gap (West Africa)"

**Table S1:** General database including geographic references and detailed information on sequenced and genotyped individuals. Empty lines indicate unsuccessful sequencing / genotyping. Haplotypes of CR1 sequences used to compute genetic diversity are provided (H1 to H14). Individuals sequenced successfully but not used to compute genetic diversity –because of missing data– are indicated by “yes”. [separate Excel file]

**Table S2:** Diversity indices calculated from the control region (CR1) among the white-bellied pangolin lineages.

*h*: Number of haplotypes, *Hd*: Haplotype (gene) diversity, *π:* Nucleotide diversity, *SD*: Standard deviation of haplotype diversity.

| Lineages | Number of individuals | *h* | *Hd* | *SD (Hd)* | *π* |
| --- | --- | --- | --- | --- | --- |
| **Dahomey Gap** | **126** | **14** | **0.686** | **0.033** | **0.00297** |
| Ghana | 3 | 2 | 0.667 | 0.314 | 0.00154 |
| Western Africa | 12 | 6 | 0.818 | 0.096 | 0.00407 |
| Western Central Africa | 59 | 31 | 0.96 | 0.012 | 0.00847 |
| Central Africa | 9 | 6 | 0.833 | 0.127 | 0.00797 |

**Table S3:** *F_ST_* values (below diagonal) and associated levels of significance (above diagonal) for the different partition schemes (i), (ii) and (iii). South and North in the partition schemes refer to Benin country.

* *p ˂ 0.05*

(i)

|  | 1 | 2 | 3 | 4 | 5 | 6 |
| --- | --- | --- | --- | --- | --- | --- |
| 1 | 0 | * |  | * | * | * |
| 2 | 0.08925 | 0 |  | * | * | * |
| 3 | 0.09822 | 0.0965 | 0 |  | * | * |
| 4 | 0.11262 | 0.16601 | 0.05282 | 0 |  | * |
| 5 | 0.10279 | 0.05366 | 0.02209 | 0.11546 | 0 | * |
| 6 | 0.24435 | 0.21227 | 0.13992 | 0.19121 | 0.13868 | 0 |

pop1 = Gnanhouizoun, pop2 = Lama, pop3 = Monts Kouffé, pop4 = Ouémé Supérieur, pop5 = Wari-Maro, pop6 = Togo.

(ii)

|  | 1 | 2 | 3 |
| --- | --- | --- | --- |
| 1 | 0.00000 | * | * |
| 2 | 0.01137 | 0.00000 | * |
| 3 | 0.03921 | 0.01865 | 0.00000 |

pop1 = South, pop2 = North, pop3 = Togo.

(iii)

|  | 1 | 2 | 3 |
| --- | --- | --- | --- |
| 1 | 0.00000 | * | * |
| 2 | 0.03783 | 0.00000 | * |
| 3 | 0.11619 | 0.06300 | 0.00000 |

pop1 = South, pop2 = North, pop3 = intermediate.

**Table S4.** Inbreeding coefficient in white-belled pangolin populations as delineated in the 6-partition scheme (i).

| **Forest** | **Fis** |
| --- | --- |
| Gnanhouizounmè | 0.098 |
| Lama | 0.098 |
| Monts Kouffé | 0.098 |
| Ouémé Supérieur | 0.131 |
| Togo | 0.317 |
| Wari-Maro | 0.101 |

**Fig S1.** Haplotype network (control region) within the Dahomey Gap lineage. Haplotype numbers refer to Table S1. Short bars correspond to mutation numbers.

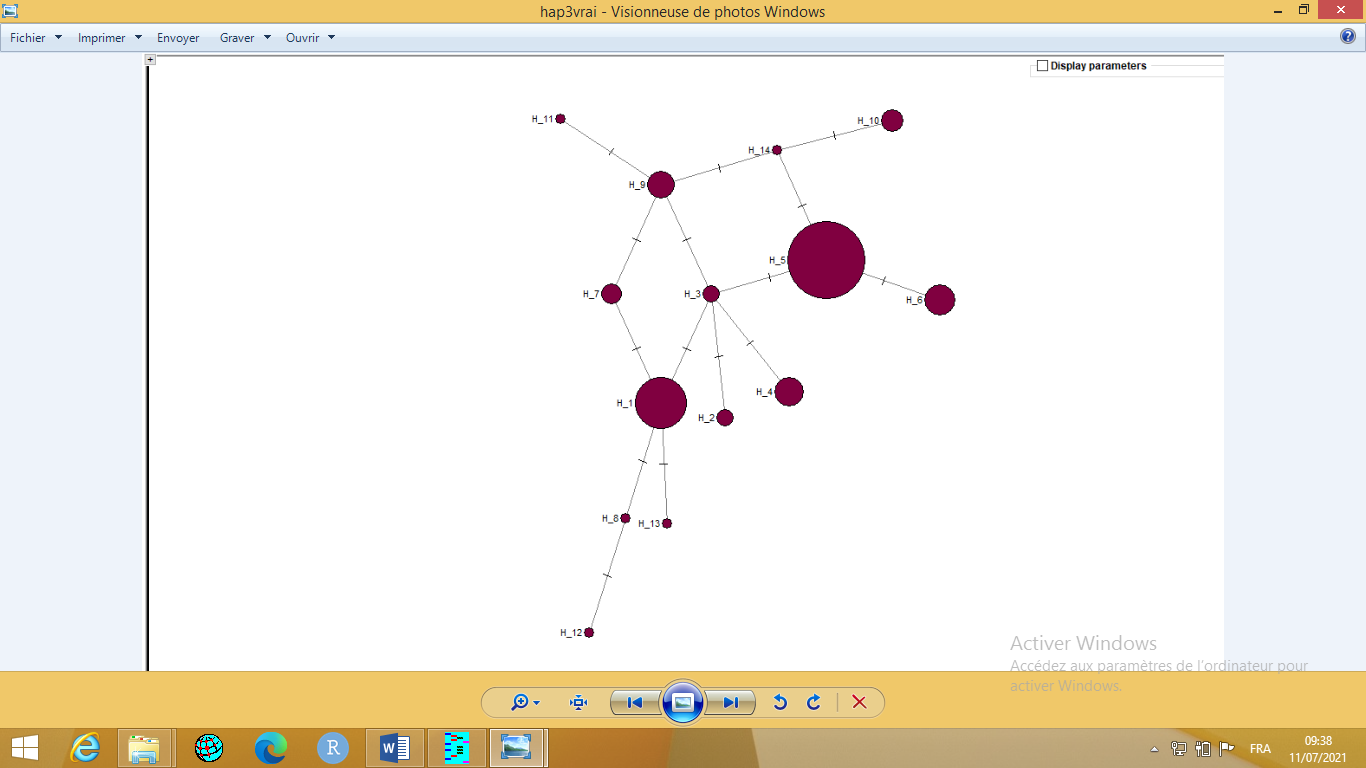

**Fig S2.** Assignment plots among the white-bellied pangolins from the Dahomey Gap as assessed with STRUCTURE for K=2 (top) and K=3 (bottom). Each individual is represented by a vertical bar.

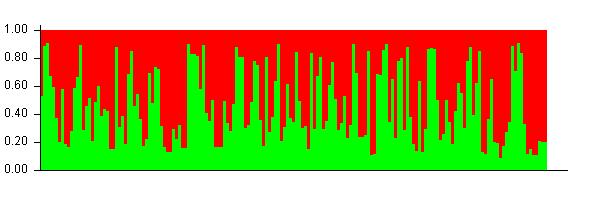

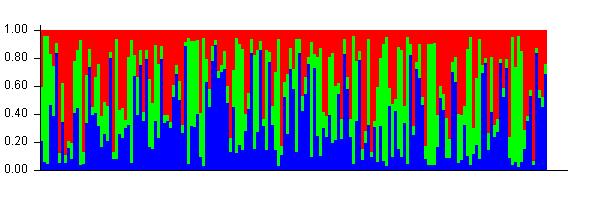

**Fig S3**. Geographic partition schemes used for computing pairwise differentiations (*F_ST_*). (i) 6-partition scheme including the forest habitats with at least 7 individuals, (ii) 3-partition scheme among South Benin-Central Benin-Togo, (iii) gradient-based 3-partition scheme within Benin.

**(i)**
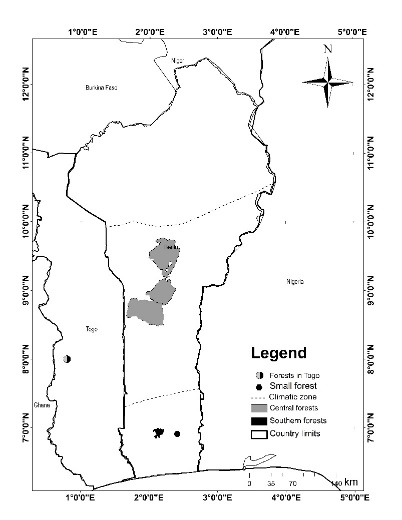

**(ii)**
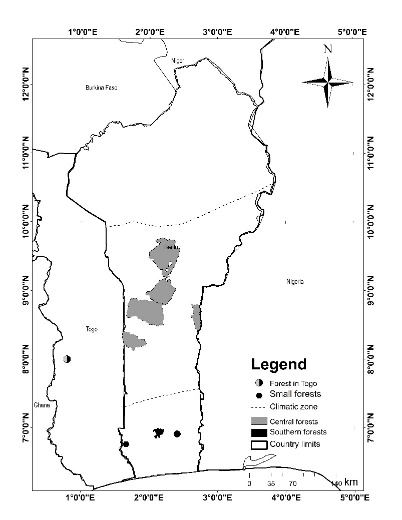

**(iii)**
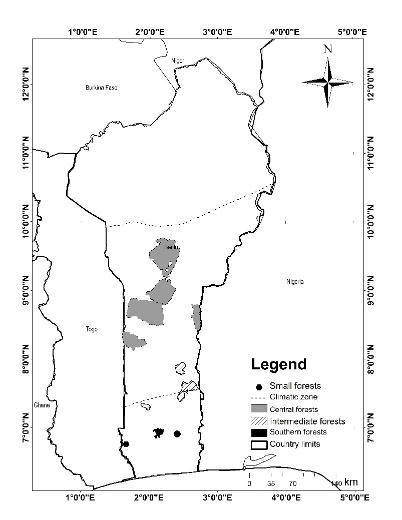

**Fig S4.** Spatial clustering of white-belled pangolins in the Dahomey Gap obtained using Geneland for K= 7 (Geneland; above) and K= 3 (Structure; below).

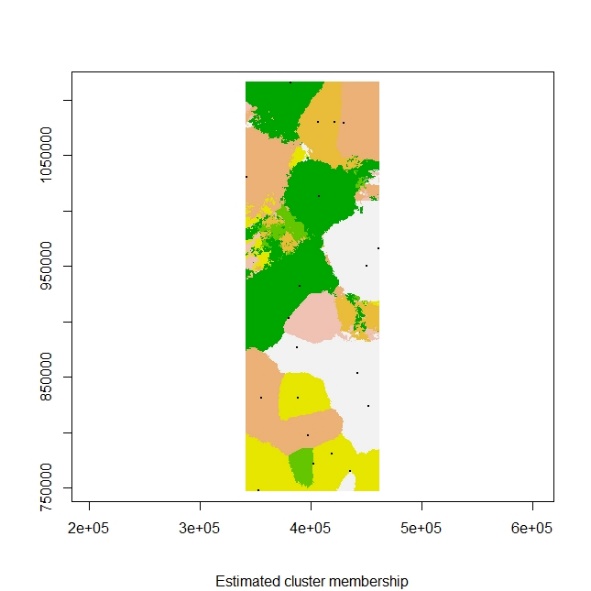

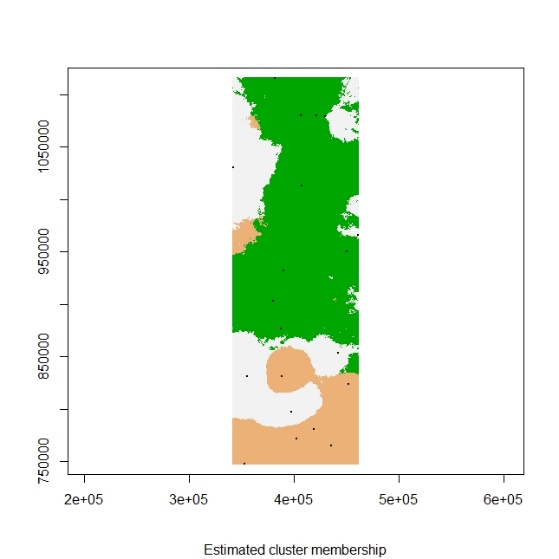

**Fig S5.** Unbiased probability of identity (uPI) and probability of identity among siblings (PIsibs) for increasing, optimized combinations among the 20 microsatellite markers.

|  | uPI | PIsibs |
| --- | --- | --- |
| 1 | 0.04124 | 0.3399 |
| 2 | 0.00299 | 0.1274 |
| 3 | 0.0002588 | 0.04965 |
| 4 | 2.973e-05 | 0.02082 |
| 5 | 3.765e-06 | 0.009128 |
| 6 | 4.778e-07 | 0.004024 |
| 7 | 7.244e-08 | 0.001786 |
| 8 | 1.388e-08 | 0.0008495 |
| 9 | 3.276e-09 | 0.0004333 |
| 10 | 7.85e-10 | 0.0002213 |
| 11 | 2.137e-10 | 0.0001263 |
| 12 | 6.329e-11 | 7.298e-05 |
| 13 | 1.953e-11 | 4.241e-05 |
| 14 | 6.176e-12 | 2.482e-05 |
| 15 | 2.875e-12 | 1.705e-05 |
| 16 | 1.936e-12 | 1.408e-05 |
| 17 | 1.451e-12 | 1.224e-05 |
| 18 | 1.112e-12 | 1.075e-05 |
| 19 | 9.38e-13 | 9.894e-06 |
| 20 | 8.124e-13 | 9.224e-06 |

**Fig S6.** Mean private allelic richness per locus in white-bellied pangolin populations (6-partition scheme), as estimated from ADZE (N=7). FGN: Gnanhouizounmè, FL: Lama, FMK: Mont Koufé, FOS: Ouémé Supérieur, FWM: Wari Maro and TG: forests of central Togo.

|  | FGN | FL | FMK | FOS | FWM | TG |
| --- | --- | --- | --- | --- | --- | --- |
| PT116 | 3.34108e-09 | 0.00327753 | 0.00333715 | 0.00821635 | 0.00621989 | 0 |
| PT175 | 0.235438 | 0.72985 | 0.2929 | 1.01902 | 0.513154 | 1.29254 |
| PT79 | 0.000756554 | 0.000122213 | 0.000600898 | 0.000204402 | 0 | 0.0214105 |
| PT197 | 0.136574 | 0.0362153 | 0.0873227 | 0.0507876 | 0.52169 | 0.437863 |
| PT839 | 0.0546158 | 0.0348604 | 0.0496397 | 0.04138 | 0.0834648 | 0.821039 |
| PT464 | 0.544282 | 0.000271945 | 0.425853 | 0.614878 | 0.524627 | 0.194991 |
| PT145 | 4.5279e-10 | 0.0527479 | 0.0206299 | 0.307984 | 0.021714 | 0.0057014 |
| PT344 | 0.15 | 0 | 0 | 0 | 0 | 0.35 |
| PT159 | 0.0153652 | 0.169726 | 0.00798312 | 7.35934e-07 | 0.0119733 | 2.32494e-05 |
| PT308 | 0.0628275 | 0.13113 | 0.32536 | 0.496547 | 0.618739 | 0.00214703 |
| PT166 | 2.4612e-06 | 2.20476e-06 | 1.95483e-06 | 2.42356e-05 | 4.18404e-06 | 6.9652e-05 |
| PT122 | 0.5 | 0 | 0.189299 | 0.0817006 | 0.172479 | 0 |
| PT739 | 0.244013 | 0.105113 | 0.0215335 | 5.70768e-07 | 0.437502 | 1.49989e-05 |
| PT619 | 0 | 0.00428077 | 0.00367993 | 0 | 0.00231875 | 0.100369 |
| PT338 | 0.452827 | 0.546177 | 0.127483 | 0.1005 | 0.150782 | 0.149632 |
| PT184 | 0.00829907 | 0.0746348 | 0.0245952 | 0.0228395 | 0.100377 | 0.0270041 |
| PT378 | 0 | 0 | 0.0197858 | 0.36796 | 0 | 0.0826018 |
| PT353 | 0.226709 | 0.322045 | 0.19634 | 0.134311 | 0.201023 | 0.4779 |
| PT276 | 2.39167e-07 | 0.749123 | 0.0263444 | 0.0497365 | 1.16103e-06 | 0.270566 |
| PT201 | 0.296568 | 0.00291798 | 0.193469 | 0.454974 | 0.0645857 | 0.549402 |

**Fig S7.** Mean private allelic frequencies per population (forest habitat), as estimated by rarefaction from ADZE. FGN: Gnanhouizounmè, FL: Lama, FMK: Mont Koufé, FOS: Ouémé Supérieur, FWM: Wari Maro and TG: forests of central Togo. The names of the three loci cross-validated (ADZE and GenAlEx) and used for the tracing (i.e. private alleles found in market individuals) appear in green. The four loci cross-validated (ADZE and GenAlEx) but not used for the tracing (i.e. private alleles not found in market individuals) appear in blue. The other four loci revealed using rarefaction but not observed in the actual dataset (GenAlEx) appear in orange.

PT_338821

|  | FGN | FL | FMK | FOS | FW | FTG |
| --- | --- | --- | --- | --- | --- | --- |
| 0 | 0% | 0% | 0% | 0% | 0% | 0% |
| 2 | 48.6215% | 44.3012% | 24.3151% | 19.7664% | 26.4337% | 28.5458% |
| 3 | 54.1296% | 52.1574% | 22.6401% | 18.7028% | 24.9607% | 23.7572% |
| 4 | 55.7415% | 56.5447% | 20.5967% | 17.2097% | 23.0076% | 21.7365% |
| 5 | 54.3592% | 57.955% | 18.2113% | 15.1361% | 20.6782% | 19.9597% |
| 6 | 50.661% | 57.0949% | 15.5537% | 12.6723% | 18.0068% | 17.692% |
| 7 | 45.2827% | 54.6177% | 12.7483% | 10.05% | 15.0782% | 14.9632% |

PT_619913

|  | FGN | FL | FMK | FOS | FW | FTG |
| --- | --- | --- | --- | --- | --- | --- |
| 0 | 0% | 0% | 0% | 0% | 0% | 0% |
| 2 | 0% | 6.25208% | 6.44121% | 0% | 3.69331% | 30.5538% |
| 3 | 0% | 4.86327% | 4.88757% | 0% | 2.82652% | 29.3215% |
| 4 | 0% | 3.19279% | 3.11542% | 0% | 1.82445% | 24.713% |
| 5 | 0% | 1.84556% | 1.73872% | 0% | 1.03628% | 19.2748% |
| 6 | 0% | 0.947455% | 0.856081% | 0% | 0.522536% | 14.2246% |
| 7 | 0% | 0.428077% | 0.367993% | 0% | 0.231875% | 10.0369% |

PT_18497228

|  | FGN | FL | FMK | FOS | FW | FTG |
| --- | --- | --- | --- | --- | --- | --- |
| 0 | 0% | 0% | 0% | 0% | 0% | 0% |
| 2 | 5.63514% | 23.6421% | 12.7774% | 11.4667% | 24.4443% | 6.7547% |
| 3 | 4.55112% | 22.6133% | 11.106% | 9.44975% | 24.357% | 6.62311% |
| 4 | 3.38836% | 19.202% | 8.64987% | 7.42203% | 21.5586% | 5.96117% |
| 5 | 2.29649% | 14.9666% | 6.13761% | 5.38138% | 17.6497% | 4.90945% |
| 6 | 1.43454% | 10.8992% | 4.02889% | 3.62782% | 13.6384% | 3.76218% |
| 7 | 0.829907% | 7.46348% | 2.45952% | 2.28395% | 10.0377% | 2.70041% |

PT_378852

|  | FGN | FL | FMK | FOS | FW | FTG |
| --- | --- | --- | --- | --- | --- | --- |
| 0 | 0% | 0% | 0% | 0% | 0% | 0% |
| 2 | 0% | 0% | 2.78517% | 26.0038% | 0% | 8.20719% |
| 3 | 0% | 0% | 3.15523% | 33.04% | 0% | 9.83264% |
| 4 | 0% | 0% | 3.12496% | 37.0746% | 0% | 10.3573% |
| 5 | 0% | 0% | 2.84739% | 38.7137% | 0% | 10.1047% |
| 6 | 0% | 0% | 2.43753% | 38.4748% | 0% | 9.33647% |
| 7 | 0% | 0% | 1.97858% | 36.796% | 0% | 8.26018% |

PT_739516

|  | FGN | FL | FMK | FOS | FW | FTG |
| --- | --- | --- | --- | --- | --- | --- |
| 0 | 0% | 0% | 0% | 0% | 0% | 0% |
| 2 | 21.3729% | 13.6458% | 6.20611% | 1.84322% | 14.9184% | 4.77455% |
| 3 | 25.579% | 15.5166% | 4.89564% | 0.40106% | 19.3204% | 1.35023% |
| 4 | 27.651% | 15.7791% | 4.28366% | 0.0666861% | 25.1054% | 0.313803% |
| 5 | 27.8488% | 14.7562% | 3.64583% | 0.00861133% | 31.2656% | 0.0622531% |
| 6 | 26.6199% | 12.8712% | 2.90684% | 0.000837731% | 37.5018% | 0.0105571% |
| 7 | 24.4013% | 10.5113% | 2.15335% | 5.70768e-05% | 43.7502% | 0.00149989% |

PT_353755

|  | FGN | FL | FMK | FOS | FW | FTG |
| --- | --- | --- | --- | --- | --- | --- |
| 0 | 0% | 0% | 0% | 0% | 0% | 0% |
| 2 | 56.7794% | 48.1656% | 54.594% | 41.8259% | 42.2066% | 35.6828% |
| 3 | 53.8178% | 47.5483% | 51.7946% | 36.7189% | 40.2704% | 34.6344% |
| 4 | 46.7821% | 44.9059% | 44.6905% | 30.4759% | 36.5993% | 36.625% |
| 5 | 38.3902% | 41.0899% | 35.9429% | 24.1776% | 31.5278% | 39.7901% |
| 6 | 30.1041% | 36.7147% | 27.273% | 18.4018% | 25.7985% | 43.5429% |
| 7 | 22.6709% | 32.2045% | 19.634% | 13.4311% | 20.1023% | 47.79% |

PT_276641

|  | FGN | FL | FMK | FOS | FW | FTG |
| --- | --- | --- | --- | --- | --- | --- |
| 0 | 0% | 0% | 0% | 0% | 0% | 0% |
| 2 | 3.76474% | 31.4562% | 9.3056% | 13.3401% | 4.78344% | 26.9262% |
| 3 | 0.59153% | 40.8145% | 7.2629% | 11.4079% | 0.896454% | 29.3888% |
| 4 | 0.070819% | 50.9299% | 6.35582% | 10.419% | 0.135466% | 31.1277% |
| 5 | 0.00658558% | 60.0921% | 5.21226% | 8.91104% | 0.0167784% | 31.0864% |
| 6 | 0.000465468% | 68.0704% | 3.90214% | 6.99188% | 0.0016403% | 29.5697% |
| 7 | 2.39167e-05% | 74.9123% | 2.63444% | 4.97365% | 0.000116103% | 27.0566% |

PT_2019332

|  | FGN | FL | FMK | FOS | FW | FTG |
| --- | --- | --- | --- | --- | --- | --- |
| 0 | 0% | 0% | 0% | 0% | 0% | 0% |
| 2 | 29.9646% | 9.39762% | 25.5224% | 45.2992% | 17.3962% | 36.2958% |
| 3 | 30.3604% | 6.24916% | 25.2624% | 46.6335% | 15.872% | 43.5929% |
| 4 | 29.6009% | 3.49502% | 23.716% | 45.8264% | 13.275% | 48.1386% |
| 5 | 29.1707% | 1.71775% | 22.1358% | 45.1359% | 10.6926% | 51.1855% |
| 6 | 29.2625% | 0.751078% | 20.7197% | 45.0541% | 8.41182% | 53.343% |
| 7 | 29.6568% | 0.291798% | 19.3469% | 45.4974% | 6.45857% | 54.9402% |

PT_1225378

|  | FGN | FL | FMK | FOS | FW | FTG |
| --- | --- | --- | --- | --- | --- | --- |
| 0 | 0% | 0% | 0% | 0% | 0% | 0% |
| 2 | 14.2857% | 0% | 12.6844% | 5.67373% | 9.7264% | 0% |
| 3 | 21.4286% | 0% | 16.3366% | 7.24414% | 12.8166% | 0% |
| 4 | 28.5714% | 0% | 18.5599% | 8.16393% | 14.9672% | 0% |
| 5 | 35.7143% | 0% | 19.5939% | 8.55595% | 16.3341% | 0% |
| 6 | 42.8571% | 0% | 19.6531% | 8.52714% | 17.0543% | 0% |
| 7 | 50% | 0% | 18.9299% | 8.17006% | 17.2479% | 0% |

PT_1594892

|  | FGN | FL | FMK | FOS | FW | FTG |
| --- | --- | --- | --- | --- | --- | --- |
| 0 | 0% | 0% | 0% | 0% | 0% | 0% |
| 2 | 7.36631% | 31.7659% | 5.92606% | 1.26842% | 10.5987% | 5.32163% |
| 3 | 6.14472% | 32.7171% | 4.4155% | 0.315636% | 6.6518% | 1.71084% |
| 4 | 4.83784% | 30.5551% | 3.20934% | 0.0596398% | 4.42395% | 0.441471% |
| 5 | 3.55207% | 26.5646% | 2.19169% | 0.00870099% | 2.99341% | 0.0943984% |
| 6 | 2.42406% | 21.7907% | 1.38192% | 0.000953541% | 1.95505% | 0.016609% |
| 7 | 1.53652% | 16.9726% | 0.798312% | 7.35934e-05% | 1.19733% | 0.00232494% |

PT_308752

|  | FGN | FL | FMK | FOS | FW | FTG |
| --- | --- | --- | --- | --- | --- | --- |
| 0 | 0% | 0% | 0% | 0% | 0% | 0% |
| 2 | 23.6958% | 40.8252% | 33.5823% | 38.2514% | 55.0495% | 9.08254% |
| 3 | 19.3636% | 37.8348% | 33.8108% | 40.7766% | 60.3736% | 5.00238% |
| 4 | 15.088% | 31.9369% | 33.5714% | 43.0859% | 62.5823% | 2.55231% |
| 5 | 11.4971% | 25.1959% | 33.3723% | 45.4692% | 63.2106% | 1.21196% |
| 6 | 8.59735% | 18.7219% | 33.0916% | 47.72% | 62.8727% | 0.533474% |
| 7 | 6.28275% | 13.113% | 32.536% | 49.6547% | 61.8739% | 0.214703% |

PT_1669238

|  | FGN | FL | FMK | FOS | FW | FTG |
| --- | --- | --- | --- | --- | --- | --- |
| 0 | 0% | 0% | 0% | 0% | 0% | 0% |
| 2 | 1.40269% | 1.47034% | 1.59221% | 5.14617% | 1.99396% | 6.17178% |
| 3 | 0.393608% | 0.406793% | 0.430541% | 1.69792% | 0.576115% | 2.20494% |
| 4 | 0.0868123% | 0.0879581% | 0.0903712% | 0.448533% | 0.131248% | 0.651092% |
| 5 | 0.0154973% | 0.0152713% | 0.0151049% | 0.0976863% | 0.0242834% | 0.16568% |
| 6 | 0.00221897% | 0.00210246% | 0.00198149% | 0.0173601% | 0.00361609% | 0.0366212% |
| 7 | 0.00024612% | 0.000220476% | 0.000195483% | 0.00242356% | 0.000418404% | 0.0069652% |

PT_1453906

|  | FGN | FL | FMK | FOS | FW | FTG |
| --- | --- | --- | --- | --- | --- | --- |
| 0 | 0% | 0% | 0% | 0% | 0% | 0% |
| 2 | 0.408016% | 26.0938% | 14.4719% | 38.083% | 10.6766% | 7.9074% |
| 3 | 0.0246442% | 22.7886% | 10.5866% | 42.335% | 8.31413% | 5.78313% |
| 4 | 0.00136134% | 17.6521% | 7.62008% | 42.2137% | 6.38888% | 3.76316% |
| 5 | 6.26309e-05% | 12.5746% | 5.18957% | 39.512% | 4.66373% | 2.20363% |
| 6 | 2.1232e-06% | 8.38735% | 3.35046% | 35.4506% | 3.24749% | 1.174% |
| 7 | 4.53e-08% | 5.27479% | 2.06299% | 30.7984% | 2.1714% | 0.57014% |

PT_464918

|  | FGN | FL | FMK | FOS | FW | FTG |
| --- | --- | --- | --- | --- | --- | --- |
| 0 | 0% | 0% | 0% | 0% | 0% | 0% |
| 2 | 29.1997% | 10.8526% | 32.3756% | 38.6292% | 32.0795% | 36.9304% |
| 3 | 35.3364% | 4.13764% | 36.4946% | 44.6869% | 36.71% | 35.4873% |
| 4 | 41.9107% | 1.43394% | 40.2627% | 50.7071% | 41.7498% | 32.6522% |
| 5 | 47.4157% | 0.442363% | 42.511% | 55.5753% | 46.1508% | 28.6775% |
| 6 | 51.5742% | 0.118937% | 43.213% | 59.1421% | 49.7132% | 24.1157% |
| 7 | 54.4282% | 0.0271945% | 42.5853% | 61.4878% | 52.4627% | 19.4991% |

PT_1973508

|  | FGN | FL | FMK | FOS | FW | FTG |
| --- | --- | --- | --- | --- | --- | --- |
| 0 | 0% | 0% | 0% | 0% | 0% | 0% |
| 2 | 25.0007% | 18.6537% | 26.2936% | 13.6534% | 31.0889% | 19.2384% |
| 3 | 26.082% | 14.2945% | 24.5441% | 13.2536% | 36.181% | 21.9891% |
| 4 | 24.3365% | 10.5002% | 20.9297% | 11.6085% | 40.4658% | 26.2953% |
| 5 | 21.1453% | 7.59614% | 16.6892% | 9.43342% | 44.4311% | 31.6996% |
| 6 | 17.394% | 5.3586% | 12.4824% | 7.16323% | 48.2725% | 37.6368% |
| 7 | 13.6574% | 3.62153% | 8.73227% | 5.07876% | 52.169% | 43.7863% |

PT_839522

|  | FGN | FL | FMK | FOS | FW | FTG |
| --- | --- | --- | --- | --- | --- | --- |
| 0 | 0% | 0% | 0% | 0% | 0% | 0% |
| 2 | 37.6771% | 28.2168% | 30.7575% | 21.0306% | 36.9956% | 44.1838% |
| 3 | 29.2942% | 20.004% | 24.4063% | 16.3048% | 31.1321% | 53.7461% |
| 4 | 20.785% | 13.3453% | 17.9267% | 12.5939% | 24.2282% | 63.0588% |
| 5 | 13.8892% | 8.61106% | 12.3317% | 9.24239% | 17.7924% | 70.7872% |
| 6 | 8.8789% | 5.4877% | 8.0267% | 6.3832% | 12.46% | 77.0224% |
| 7 | 5.46158% | 3.48604% | 4.96397% | 4.138% | 8.34648% | 82.1039% |

PT_1162028

|  | FGN | FL | FMK | FOS | FW | FTG |
| --- | --- | --- | --- | --- | --- | --- |
| 0 | 0% | 0% | 0% | 0% | 0% | 0% |
| 2 | 1.02253% | 8.49556% | 8.9595% | 13.3138% | 10.1237% | 0.32727% |
| 3 | 0.0975024% | 5.15584% | 5.90987% | 9.56132% | 7.15315% | 0.0155865% |
| 4 | 0.00704221% | 2.97111% | 3.39297% | 5.96855% | 4.42266% | 0.000511911% |
| 5 | 0.000380246% | 1.56034% | 1.73552% | 3.36024% | 2.48066% | 9.8825e-06% |
| 6 | 1.44042e-05% | 0.748058% | 0.801002% | 1.73336% | 1.2866% | 7.9e-08% |
| 7 | 3.341e-07% | 0.327753% | 0.333715% | 0.821635% | 0.621989% | 0% |

PT_1753627

|  | FGN | FL | FMK | FOS | FW | FTG |
| --- | --- | --- | --- | --- | --- | --- |
| 0 | 0% | 0% | 0% | 0% | 0% | 0% |
| 2 | 23.6406% | 43.5772% | 29.5642% | 63.3342% | 38.3827% | 80.3803% |
| 3 | 24.6297% | 52.1627% | 33.9164% | 80.5775% | 47.3198% | 104.514% |
| 4 | 25.8989% | 60.6021% | 35.36% | 91.7719% | 52.7698% | 120.266% |
| 5 | 26.155% | 67.0307% | 34.5323% | 98.2504% | 54.8306% | 128.843% |
| 6 | 25.2995% | 71.096% | 32.2887% | 101.281% | 54.1333% | 131.467% |
| 7 | 23.5438% | 72.985% | 29.29% | 101.902% | 51.3154% | 129.254% |

PT_796077

|  | FGN | FL | FMK | FOS | FW | FTG |
| --- | --- | --- | --- | --- | --- | --- |
| 0 | 0% | 0% | 0% | 0% | 0% | 0% |
| 2 | 5.50645% | 1.61522% | 6.25047% | 2.58085% | 0% | 24.2284% |
| 3 | 3.26891% | 0.881323% | 3.57563% | 1.42126% | 0% | 18.3119% |
| 4 | 1.60674% | 0.392953% | 1.67261% | 0.639612% | 0% | 12.0506% |
| 5 | 0.678883% | 0.14812% | 0.661694% | 0.24334% | 0% | 7.25787% |
| 6 | 0.246894% | 0.0470274% | 0.220472% | 0.0779666% | 0% | 4.07467% |
| 7 | 0.0756554% | 0.0122213% | 0.0600898% | 0.0204402% | 0% | 2.14105% |

PT_34432

|  | FGN | FL | FMK | FOS | FW | FTG |
| --- | --- | --- | --- | --- | --- | --- |
| 0 | 0% | 0% | 0% | 0% | 0% | 0% |
| 2 | 10.8333% | 0% | 0% | 0% | 0% | 20.7143% |
| 3 | 13.9286% | 0% | 0% | 0% | 0% | 27.5% |
| 4 | 15.7143% | 0% | 0% | 0% | 0% | 32.1429% |
| 5 | 16.369% | 0% | 0% | 0% | 0% | 34.8214% |
| 6 | 16.0714% | 0% | 0% | 0% | 0% | 35.7143% |
| 7 | 15% | 0% | 0% | 0% | 0% | 35% |

**Fig S8**. Neighbor joining tree inferred from the control region and including the six lineages of white-bellied pangolins. The tree includes 181 sequences from the Dahomey Gap, 59 sequences from Western Central Africa, 12 sequences from Western Africa, 9 sequences from Central Africa, 3 sequences from Ghana and 1 sequence from Gabon. [separate pdf]
