## Supplementary figures and images for "Can DNA help trace the local trade of pangolins? A genetic assessment of white-bellied pangolins from the Dahomey Gap (West Africa)"

### Supplemental Figure 8

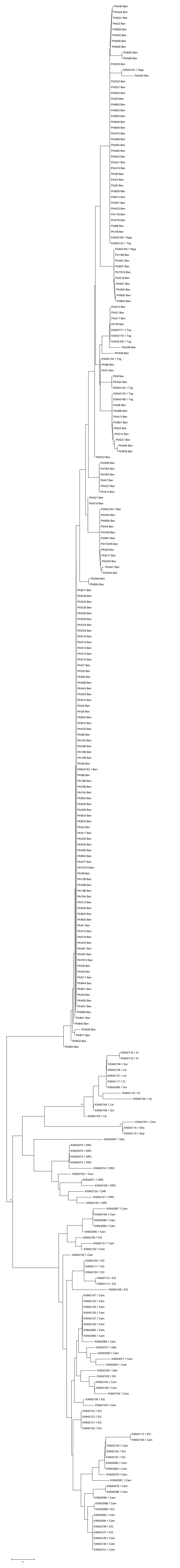
